# Structures of *Arabidopsis thaliana* MDL Proteins and Synergistic Effects with the Cytokine MIF on Human Receptors

**DOI:** 10.1101/2023.01.30.525655

**Authors:** Lukas Spiller, Ramu Manjula, Franz Leissing, Jerome Basquin, Priscila Bourilhon, Dzmitry Sinitski, Markus Brandhofer, Sophie Levecque, Björn Sabelleck, Regina Feederle, Andrew Flatley, Ralph Panstruga, Jürgen Bernhagen, Elias Lolis

## Abstract

Vertebrates have developed effective immune mechanisms to fight microbial attacks, relying on a sophisticated network of innate and adaptive responses, a circulatory system, and numerous orchestrating soluble mediators such as cytokines. Mammalian macrophage migration inhibitory factor (MIF) and its paralog D-dopachrome tautomerase (D-DT/MIF-2) are multifunctional inflammatory cytokines with chemokine-like properties that modulate immunity. Plants possess orthologous MIF/D-DT-like (MDL) proteins, whose function is largely unexplored. Driven by the previous discovery of cross-kingdom mimicry of plant (*Arabidopsis thaliana*) MDL proteins and human MIF receptor signaling, we here characterized the structures of the three *A. thaliana* MDLs by X-ray crystallography and explored the mechanism underlying the interplay between plant MDLs, human MIF, and its receptors. We obtained high-resolution structures at 1.56 Å, 1.40 Å, and 2.00 Å resolution for MDL1, MDL2, and MDL3, respectively, revealing a typical trimeric assembly and a high three-dimensional similarity to human MIF. Although residues at the catalytic site of the three MDLs show high identity to human MIF, the proteins showed low tautomerase activity for the substrate 4-hydroxyphenylpyruvate (HPP). Structural differences likely explain the enzymatic inactivity of plant MDLs for HPP. Strikingly, employing *in vitro, in vivo,* and *in planta* test systems, we found that MIF and MDL proteins interact with each other and have the capacity to form hetero-oligomeric complexes. The functional consequences of this interaction were demonstrated applying a yeast-based reporter system specific for the MIF chemokine receptors CXCR2 and CXCR4. MDLs not only triggered receptor signaling on their own, but exhibited pronounced synergism regarding the activation of the CXCR2- and CXCR4-dependent signaling pathways, when co-applied with MIF. These findings were substantiated by the co-administration of pharmacological inhibitors that either disrupt MIF receptor binding or block the catalytic cavity. Moreover, biochemical and biophysical experiments using an allosteric oligomer-specific MIF inhibitor established hexa-oligomer formation between MIF and MDLs as the putative basis for the synergistic effect. Our results are the starting point for a mechanistic understanding of the immunomodulatory activity of a family of highly conserved plant proteins.

**One Sentence Summary:** *A. thaliana* MDLs and human macrophage inhibitory factor interact with each other and induce synergistic effects in activating CXCR2 and CXCR4.

## INTRODUCTION

The immune defense system of vertebrates relies on a sophisticated network of innate and adaptive arms and is composed of a remarkable variety of immune cells that communicate and traffic via a professional circulatory system (*1*). Cytokines and chemokines are specialized soluble immune mediators and act as coordinators of the human immune response. Accordingly, dysregulated cytokine and chemokine responses are associated with numerous diseases (*2*). Macrophage migration inhibitory factor (MIF) and its paralog D-dopachrome tautomerase (D-DT/MIF-2) are multifunctional inflammatory cytokines with chemokine-like properties that are key components of the host immune response (*3–6*). MIF not only signals through its cognate receptor CD74 to control proliferation, survival, and inflammatory responses (*7*), but also engages in non-cognate interactions with the chemokine receptors CXCR2, CXCR4, and CXCR7 to promote immune cell recruitment (*8*). These activities also causally link MIF to a variety of human diseases including acute and chronic inflammatory conditions, atherosclerosis, autoimmune disorders, neurodegenerative diseases, and cancer (*4, 8–14*).

MIF-/D-DT-like (MDL) proteins have been identified in almost all kingdoms of life, including uni- and multicellular parasites, fungi and plants, suggesting that the evolutionary origin of the ancestral MIF/D-DT gene dates back over 900 million years (*15–17*). Parasite-derived MIF orthologs can mimic mammalian MIF activities to act as virulence factors as a basis for immune evasion and are in some cases pharmacological targets (*16*). Plants have developed effective innate immune mechanisms such as pattern recognition receptors (PRRs) to fight microbial attacks but lack an adaptive immune system (*18*). Moreover, many of the primordial organisms expressing *MIF*-like genes lack a circulation and a cell-based immune system and in some, even the existence of G protein-coupled receptors (GPCRs), which act as secondary MIF receptors in vertebrates, is controversial. These facts have fueled speculation about MIF as an ancient enzyme throughout evolution, which acquired extracellular functions as a cytokine in a process of neofunctionalization (*17*).

The three-dimensional (3D) structure of human MIF (*19*) bears striking resemblance to a group of bacterial enzymes consisting of 4-oxalocrotonate tautomerase (4-OT), 5-(carboxymethyl)-2-hydroxymuconate isomerase (CHMI), and malonate semialdehyde decarboxylase (MSAD) (*20*). The MIF monomer has a molecular mass of 12.5 kDa, comprising two ⍺-helices tightly packed against four antiparallel oriented β-strands. However, MIF crystallizes as a homotrimer, in which three monomers interact with each other to form a barrel-shaped structure with a central solvent channel running through the protein assembly (*19*). Human MIF shares a sequence identity of < 20% with the above-mentioned microbial enzymes, but has a tautomerase catalytic cavity between its subunits and an unusually acidic N-terminal proline residue with a pKa of 5.6, consistent with a function as a catalytic base (*21–23*). While MIF can catalyze the tautomerization of the non-physiological substrate D-dopachrome (or D-dopachrome methyl ester [DME]) and enol-keto forms of the physiological molecule 4-hydroxyphenylpyruvate (HPP) *in vitro* (*21, 22*), a *bona fide* substrate for MIF activity in humans has remained elusive and a role of the enzymatic activity in human cells is unclear. However, mutational and inhibitor studies have demonstrated that changes of the catalytic cavity lead to conformational alterations, which affect MIF binding to CD74, CXCR2, and CXCR4 (*24–28*). The catalytic site was used by a variety of methods to identify small molecule inhibitors that affect several mouse models of disease. While much less studied, MIF also features redox-regulatory activity related to its redox-sensitive cysteine residues (*29*) and has been suggested to have nuclease activity owing to a PD-D/E(X)K nuclease motif (*30*). Comparison of MIF/MDL proteins across different kingdoms revealed a high degree of sequence conservation, with many sites being under selection in some kingdoms, especially in plants (*17, 31*). Conservation is high for the tautomerase site, while other motifs known to be of functional importance in human MIF (e.g., the pseudo-*E*LR motif required for CXCR2 binding) are not well conserved (*32*). There seems to be a complex interplay between vertebrate host proteins and its parasite MIF orthologs, with implications for virulence and host defense, but the underlying mechanisms are incompletely understood (*15, 16, 33*).

Even less understood is the relevance of MIF/D-DT-like proteins (MDLs) in the plant kingdom. *In silico* analysis revealed that MDL genes/proteins have an extraordinary degree of evolutionary conservation, but little is known about their functional roles in plants (*17, 34*). Multiple *MDL* genes are typically present per plant species, including model plants such as *Arabidopsis thaliana*, crops and other food plants. The three *A. thaliana* MDLs (*At*MDLs; herein termed MDLs for simplicity) share a sequence identity of 28–33% with human MIF, with higher conservation in the tautomerase cavity (SFig. 1) (*17*). While MDL1 and MDL2 localize in the cytoplasm of plant cells, MDL3 resides in peroxisomes (*34*). Of note, experimental evidence indicates that the tautomerase activity of MDLs for the artificial substrates HPP and D-dopachrome is dramatically reduced in comparison to human MIF (*33*). Given the sequence homology between MDLs and human MIF, we previously tested whether MDLs would interact with components of the human MIF signaling network, similar to the virulence paradigm established for parasite MIF orthologs. We observed a surprising degree of cross-kingdom mimicry, with MDLs binding to and activating the human MIF receptors CXCR4 and CD74 and promoting the chemotaxis of human leukocytes. This observation expanded the previously established interplay between the plant immune system and MIF proteins delivered by the plant-parasitic aphid *Acrythosiphon pisum* to an unanticipated cross-kingdom communication axis between components of the plant and human immune system (*35*).

In this study, we sought to characterize the structures of MDLs and understand the mechanisms underlying the interplay between plant MDLs, human MIF (herein termed MIF), and its receptors. We determined the crystallographic structures of all three *A. thaliana* MDLs, identified structural similarities between MDLs and MIF, and unraveled the basis for the surprisingly low tautomerase activity of MDLs. We finally demonstrated by biochemical, cell biological, and biophysical methodologies that MDLs and MIF form hetero-oligomeric complexes to tailor MIF-driven receptor responses by cross-kingdom synergy.

## RESULTS

### Crystal structures of the *A. thaliana* MIF orthologs MDL1, MDL2, and MDL3, and examination of their tautomerase site

We expressed and purified recombinant hexahistidine-tagged MIF orthologs MDL1-6xHis, MDL2-6xHis, and MDL3-6xHis (thereafter referred to as MDL1, MDL2, and MDL3) and solved their X-ray structures to 1.56 Å, 1.40 Å, and 2.00 Å resolution, respectively (Fig. 1A; SFig. 2; STable 1). All three MDL proteins crystallized as trimers with a very high overall structural similarity to the human MIF trimer (Fig. 1). Analysis of a structure-based alignment revealed 27% sequence identity for an all-against-all comparison of all three MDLs and 12% sequence identity for a comparison of the three MDLs with MIF (SFig. 1). The 14 invariant residues per monomer in the structural alignment (SFig. 3A) are localized into separate regions: region 1 is the catalytic cavity between two subunits and contains Pro-1 and Ser-63, while region 2 is a discontinuous surface outside the catalytic cavity composed of the six residues Ala-27, Gly-31, Pro-33, Gly-65, Ser-63, and Asp-100 (SFig. 3A,B). In region 3, a major portion of the

**Figure 1.**
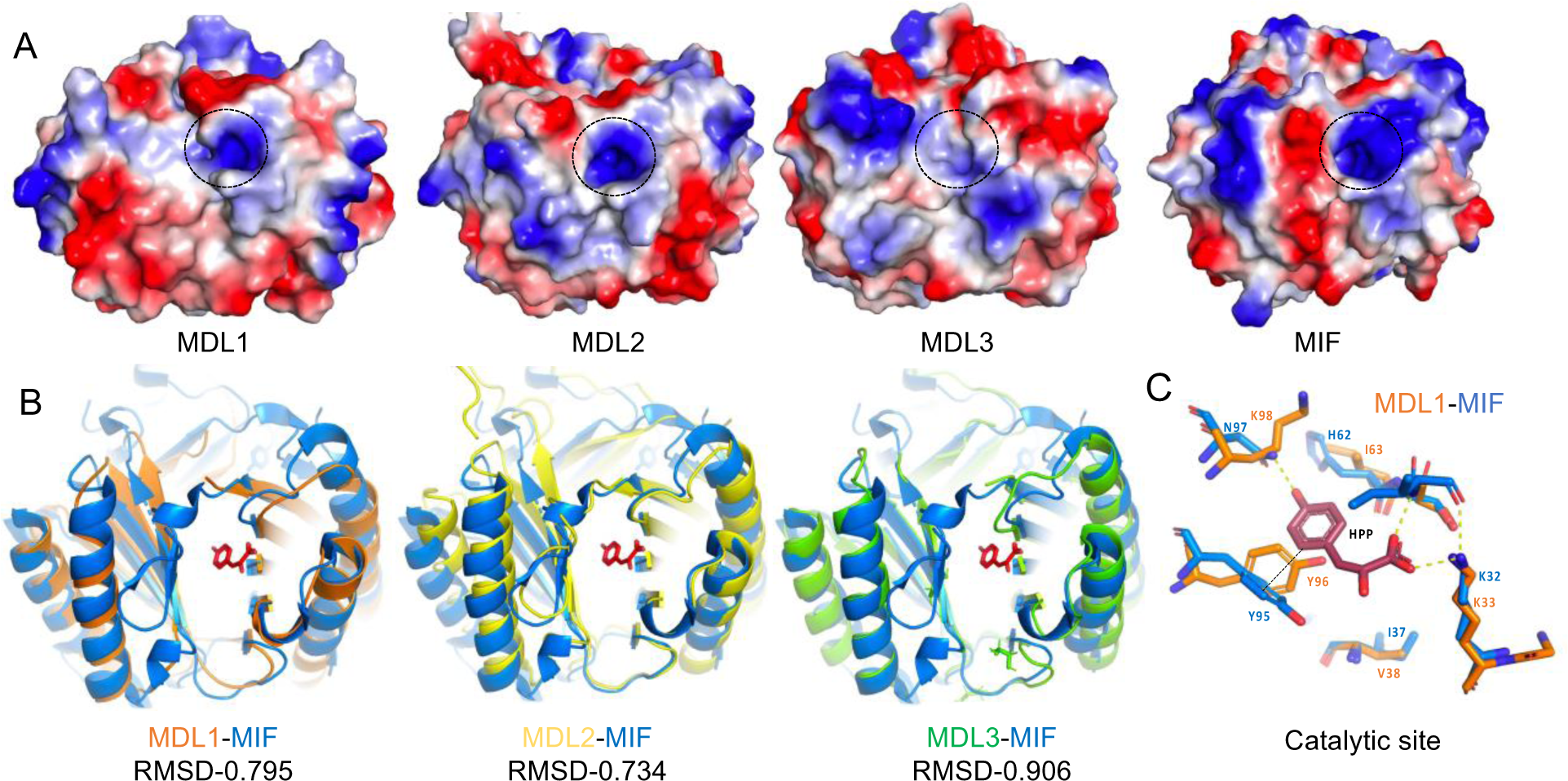
Structural properties of each MDL protein and comparison to MIF. (**A**) Electrostatic surface potential of each *A. thaliana* MDL (MDL) for substrate binding site (marked with a black circle) in comparison to human MIF (MIF). Regions of negative potential are colored red, those of positive potential are colored blue, and neutral regions are shown in white-gray. (**B**) Structural overlays of MDLs on the complex of MIF-HPP (PDB-1CA7). The blue cartoon represents the structure of MIF, orange, yellow and green cartoons represent MDL1, 2 and 3, respectively. The root-mean-square deviation (RMSD) of atomic positions is shown for each complex. The HPP in these overlays is used as the position of the modelled HPP in the MDL-HPP complexes for analysis. (C) MDL1 catalytic residues (orange Cα atoms) superimposed on MIF (Cα atoms from PDB-1CA7). The hydrophilic interactions between MIF and HPP are represented as yellow lines, and the aromatic interaction is shown as a black line.

Asp-100 surface area is outside the solvent channel, adjacent to the MIF allosteric site residue Tyr-99, whose side chain is within the channel serving as a solvent gating residue (SFig. 3C) (*26*). Interestingly, MDL1 and MDL2 also have a tyrosine residue at the equivalent position, while MDL3 has a phenylalanine residue (SFig. 1). Of note, there are three residues that belong to multiple regions based on their structural orientation. For example, Ser-63 is part of both regions 1 and 2 with the hydroxyl group being part of the catalytic cavity and its backbone contributing to the surface area (SFig. 3B). Asp-100 belongs to region 2, where Gly-65 makes a hydrogen bond between their backbone atoms, but the side chain of Asp-100 is the only residue in region 3. At region 4, the Arg-93 side chain makes a hydrogen bond to the backbone of Phe-49, which is at the C-terminal end of a β-strand involved in subunit-subunit interactions and serves a role in stabilizing this β-strand that provides MDL/MIF specificity to form homotrimers (SFig. 3D). Of the remaining five residues, Ala-57 and Leu-83 are buried within the hydrophobic core of the protein and Thr-7, Asn-8, and Gly-51 are localized in loop regions (SFig. 3A).

We also examined the catalytic cavity in more detail, focusing only on the structures of the MDLs. With the exception of MDL3, a view of the electrostatic potential of MDL1 and MDL2 does not explain the large difference in catalytic activity between MIF and MDL1 or MDL2, as they all display a positive electrostatic potential at the active site (Fig. 1A). We therefore superimposed each MDL on the MIF-HPP enzyme-substrate complex to create a model of HPP interacting with the MDLs’ cataytic site (*36*). The interactions were analyzed and compared to the respective MIF-HPP complex. The major difference in catalytic residues between human MIF (Pro-1A, Lys-32A, Ser-63A, Ile-64A, Tyr-95C, and Asn-97C; A, B, and C refer to the trimer subunits) and the three MDLs is Lys-98C (*note*: MDL Lys-98 is equivalent to MIF Asn-97 due to an extra residue in the alignment as shown in SFig. 1). Substitution of Asn-97 in MIF by lysine dramatically reduced the tautomerase activity for both HPP and DME (*33*), suggesting that Lys-98 is responsible for the lack of any robust enzymatic activity for all three MDLs using these artificial substrates. The major structural difference between MDL1 and MIF to explain the decrease of enzymatic activity is the different side chain orientation of Lys-98, which is oriented away from HPP with a distance >5.4 Å for all three subunits in MDL1. By contrast, the side chain amide group of Asn-97 in MIF forms a hydrogen bond with HPP (Fig. 1C). The large distance between Lys-98 of the MDLs and HPP is similar in the analysis for MDL2 and MDL3 with the modelled HPP, resulting in a loss of a hydrogen bond interaction and decreased affinity for HPP (SFig. 4A,B).

An unanticipated difference is observed in MDLs at residue 96, which is the equivalent position of MIF Tyr-95 (SFig. 1). The side chain of Tyr-96 for MDL1 has different conformations in the three subunits. In one subunit it clashes with the modelled HPP and in other the two subunits the side chain has no predicted interactions with HPP. The equivalent residues for MDL2 and MDL3 are Phe-96 and Ile-96, respectively. The proteins differ in the position of these residues from Tyr-96 in MDL1, with Phe-96 of MDL2 making van der Waals interactions with HPP, while the Ile-96 of MLD3 is not predicted to interact with HPP at all.

### MIF and its *A. thaliana* orthologs engage in direct protein-protein interactions *in vitro*, *in vivo*, and *in planta*

The high degree of structural similarity between MIF and the three MDLs and the capacity of each of these proteins to form homo-trimers prompted us to investigate whether these proteins would also physically interact with each other across kingdom boundaries. To test this possibility experimentally, we first performed *in vitro* co-immunoprecipitation assays with MIF and MDL1 as a representative of the three MDLs. MDL1-6xHis and biotinylated MIF-6xHis were mixed, complexes pulled down by streptavidin-coated magnetic beads, and the resulting eluate analyzed by immunoblot with 6xHis- and custom-made MDL1-specific antibodies (for MDL1 antibody validation see SFig. 5). This revealed association of the recombinant MIF and MDL1 proteins (Fig. 2A). We next asked whether the interaction between MIF and MDL proteins may also occur under *in vivo* conditions. In yeast-two-hybrid assays, we tested all pairwise interactions between MIF, MDL1, MDL2 and MDL3. Similar to our previous data (*34*), we detected a weak homomeric MDL1-MDL1 interaction and a strong heteromeric MDL1-MDL2 interaction in this system. In accordance with earlier biochemical evidence (*19, 37, 38*), we also noticed a homomeric MIF-MIF interaction. MIF-MDL2 complex formation occurred in yeast when MDL2 was used as bait protein (Fig. 2B), but not when MIF was used as bait. To substantiate these findings suggesting direct cross-kingdom binding between human MIF and a plant MDL, we performed *in planta* luciferase complementation imaging (LCI) assays. In this experimental setup, fusion proteins tagged with enzymatically inactive N- and C-terminal segments of firefly luciferase (nLUC and cLUC, respectively) are transiently expressed in plant leaves. Interaction of candidate proteins results in the reconstitution of enzymatically active luciferase, which can be detected and quantified upon addition of a suitable substrate (e.g., luciferin). Co-expression of nLUC-MIF with cLUC-tagged MDL1, MDL2, MDL3 or MIF resulted in strong luciferase activity for the cLUC-MDL2 and nLUC-MIF combination. Similarly, expression of cLUC-MIF yielded strong luciferase activity in the reciprocal combination with nLUC-MDL2 and additionally with nLUC-MIF (Fig. 2C, D; SFig. 6). To quantify direct binding between MIF and MDL homologues, we determined the KD values of MIF-MDL interactions using microscale thermophoresis (MST), a biomolecular interaction methodology suitable to measure protein-protein binding at nano- to micromolar concentrations under solution conditions. We chemically labeled recombinant MIF with the RED-NHS dye to analyze the interaction with unlabeled recombinant MDL1, MDL2, MDL3, respectively. We observed characteristic sigmoidal binding curves with KD values less than 5 µM for each MIF-MDL pair (Fig. 2E). Negative controls did not result in sigmoidal binding curves, indicating the MIF-MDL interactions were due to specific binding (SFig. 7). Taken together, four different types of protein-protein interaction assays (*in vitro* co-immunoprecipitation and MST, *in vivo* yeast two-hybrid, and i*n planta* LCI experiments) provided evidence for direct association of MIF and MDL proteins.

**Figure 2.**
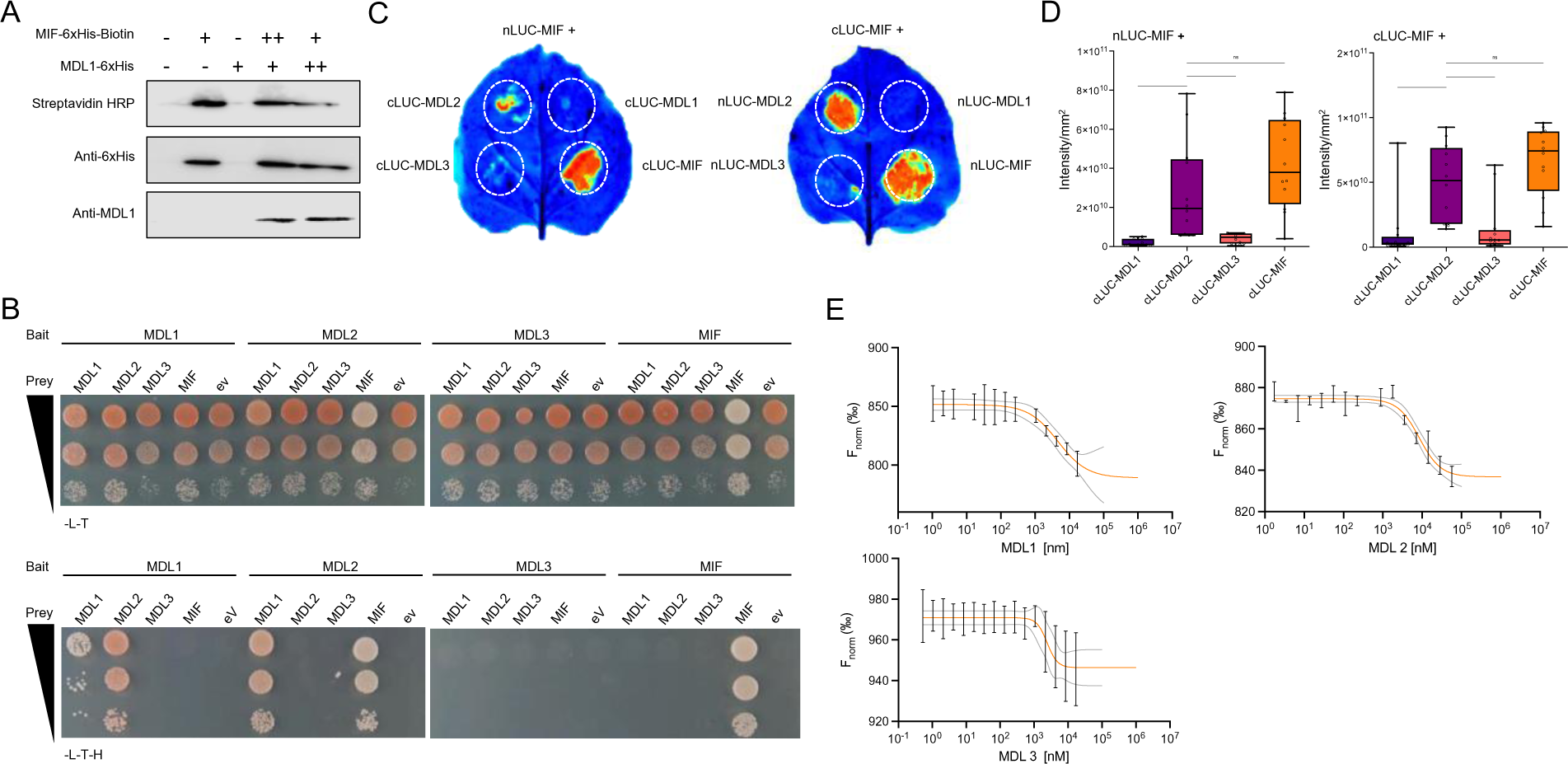
*In vitro, in vivo,* and *in planta* interaction between MIF and MDL proteins. (**A**) Co-immunoprecipitation of MIF and MDL1. Biotin-MIF-6xHis and -MDL1-6xHis were co-incubated at indicated ratios and complexes pulled down by streptavidin-coated beads (–, protein not present; +, 500 ng protein added; ++, 1000 ng protein added). (**B**) Interaction between MIF and MDL proteins in a yeast two-hybrid assay. All possible bait/prey combinations were tested as indicated. Growth control (upper panel) was performed on synthetic complete medium lacking leucine (-L, selection for bait vector) and tryptophan (-T, selection for prey vector). Selection for interaction (lower panel) was performed on synthetic complete medium lacking leucine (-L), tryptophan (-T) and histidine (-H, selection for interaction); ev, empty vector. Shown is each in panel from top to bottom for each construct a 10-times dilution series. Photographs were taken after 3 d of yeast growth. Two additional experimental replicates yielded a similar outcome. (**C**) and (**D**) *In planta* interaction between MIF and MDL proteins as determined in a luciferase complementation imaging assay. (**C**) Luminescence images of representative leaves are shown. The circles marked by dashed white lines indicate the approximate agrobacteria infiltration sites. Warmer colors indicate a higher level of luminescence. (**D**) Quantification of the luminescence signals according to (**C**). The intensity of light emission was measured and calculated per square mm. The experiment was performed three times with four leaves each. Boxplots show the results of the 12 data points per combination. (**E**) Direct protein–protein interaction studies between fluorescently labeled RED-NHS-MIF and MDL proteins using microscale thermophoresis (MST). For a constant MIF concentration of 100 nM, the difference in normalized fluorescence [‰] is plotted against increasing MDL concentrations for analysis of thermophoresis. Values shown represent means ± SD as obtained from at least three independent experiments. Further control experiments are shown in SFig. 7.

### Synergy by MIF and MDLs on human chemokine receptor activation

We have previously used a modified genetic strain of *S. cerevisiae* that expresses functional human chemokine receptors, which can be activated by chemokines and signal through heterotrimeric proteins using an altered *S. cerevisiae* Gα (GPA1) subunit. The signaling pathway downstream of GPA1 activation includes the mitogen-activated protein kinase (MAPK) pathway, the STE12 transcription factor, and substituted genes transcribed by STE12, which allow the β-galactosidase (*lacZ/ß-gal*) reporter system to function (*25, 39–42*) (Fig. 3A). Capitalizing on this established system for CXCR4, we tested MIF and the MDLs for activation and signaling of CXCR4 and CXCR2 (*25*) (Fig. 3B). Interestingly, both MDL1 and MDL2 activated CXCR4 more potently than MIF, with each protein used at 20 µM. When 10 µM MIF and 10 µM of either MDL1 or MDL2 were tested together, a hyper-activated (synergistic) effect was observed, with the MIF-MDL2 mixture approximately three times more active than the MIF-MDL1 combination (Fig. 3B). We explored the dose-dependent responses of MIF from 1-20 µM alone and 20 µM MIF in the presence of 1-4 µM MDL1. While there was no difference in activation with different concentration of MIF alone in this dose range, there was an increasing level of synergy, even when MIF was combined with lower concentrations of MDL1 (Fig. 3C).

**Figure 3.**
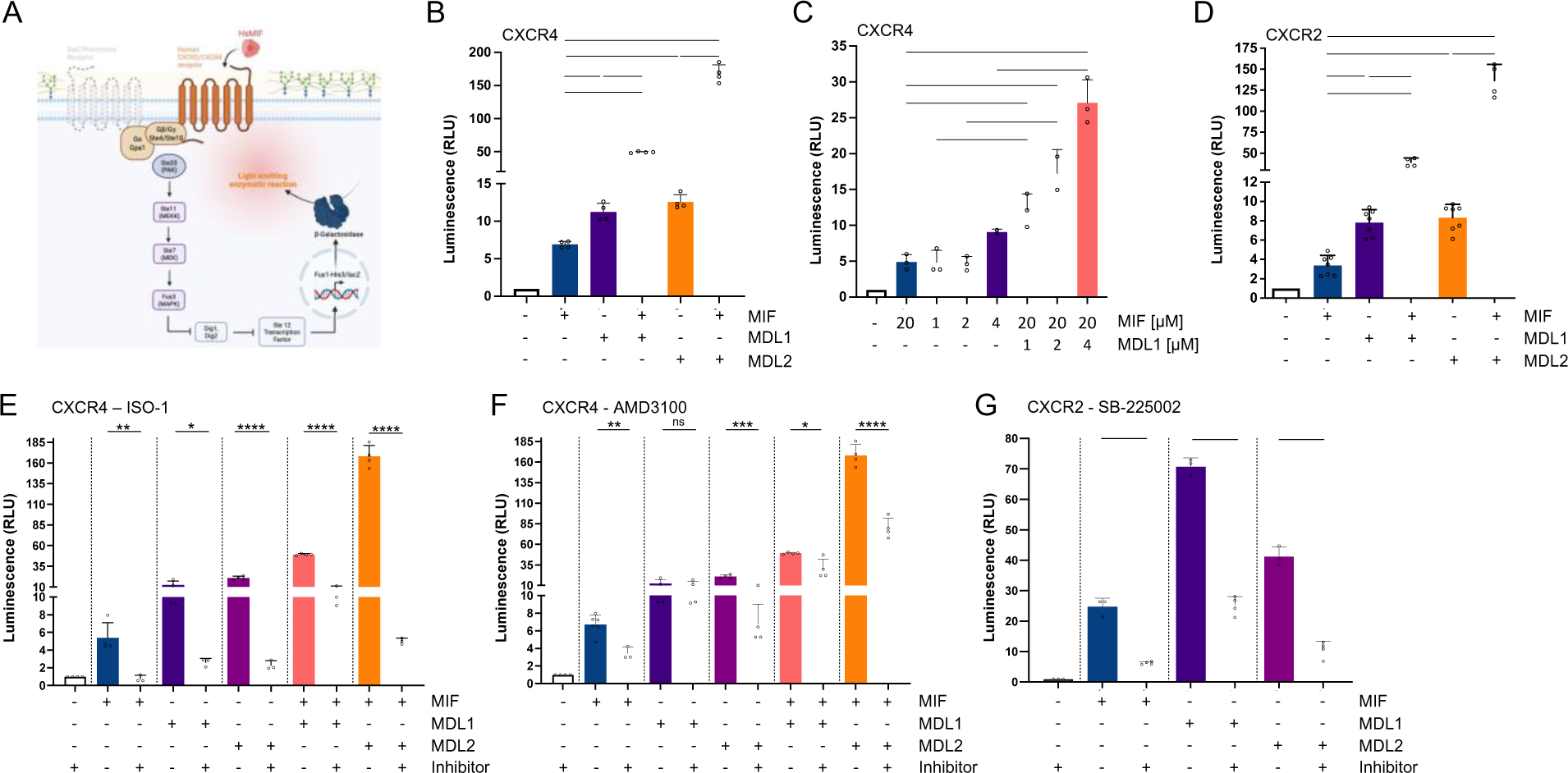
Synergistic effect of MIF and MDL proteins on human chemokine receptor activation in a yeast-based reporter system. CXCR2 and CXCR4 receptor binding and signaling activation in a yeast-based reporter system following addition of MIF and MDL recombinant proteins, alone or in combination in the absence or presence of specific inhibitors. (**A**) Schematic illustration of the modified pheromone-response signaling pathway in *S. cerevisiae.* The endogenous Ste2 GPCR has been replaced by the human chemokine receptors CXCR2 or CXCR4, respectively and linked to the downstream signaling cascade. Ligand binding results in activation of the MAP-kinase like pathway and eventually triggers expression of the *lacZ* reporter gene. Resulting β-galactosidase activity was measured photometrically upon addition of the corresponding substrate. (**B**) Luminescence (in relative light units, RLU) due to *lacZ* reporter gene activation in the CXCR4 yeast system after addition of recombinant proteins. MIF and MDLs were used either individually or in different 1:1 combinations, at a final total concentration of 20 µM protein per treatment. (**C**) Titration experiment in a yeast-based CXCR4 reporter system. Increasing concentrations (1, 2, 4 µM) of MIF alone or in combination with 20 µM MDL1 were added. For comparison, the effect of MIF at a concentration of 20 µM is shown. (**D**) Luminescence (in RLU) due to *lacZ* reporter gene activation in the CXCR2 yeast system after addition of recombinant proteins. MIF and MDLs were used either individually or in different 1:1 combinations, at a final total concentration of 20 µM protein per treatment. (**E-G**) Effects of known MIF or chemokine receptor inhibitors on MIF/MDL-mediated receptor activation in yeast-based CXCR2/CXCR4 reporter systems. The plots show luminescence (in RLU) due to *lacZ* reporter gene activation upon the addition of MIF and MDL recombinant proteins alone or in 1:1 combinations, at a final total protein concentration 20 µM, either in the absence or presence of 100 µM ISO-1 (CXCR4 reporter system (**E**); 100 µM AMD3100 in the CXCR4 reporter system (**F**); or 200 µM SB225002 in the CXCR2 reporter system (**G**)). Values shown represent means ± SD as obtained from at least three independent experiments with RLUs of each experiment assessed in technical duplicates and normalized to untreated controls. Individual data points are indicated by white circles. Statistical analysis was performed using one-way ANOVA with multiple comparison (* p < 0.05, ** p < 0.01, *** p < 0.001, **** p < 0.0001).

The activation of chemokine receptor CXCR2 by MIF has been shown in mammalian cells (*8, 28, 32*), but this pathway is also activated by MIF in *S. cerevisiae* in a reporter system analogous to the one described above for CXCR4 (Fig. 3D). MDLs lack the pseudo-*E*LR motif of two non-adjacent residues, Arg-11 and Asp-44, present in human MIF, which contributes to binding and activation of CXCR2 (*32*). Therefore, MDL1 and MDL2 were not expected to activate CXCR2. However, application of 20 µM MIF, MDL1, and MDL2 revealed that MDL1 and MDL2 activate CXCR2 to a greater extent than MIF, even though the MDL proteins contain uncharged residues in positions 11 and 44 instead of charged residues (Fig. 3D, SFig. 1). Given the results with CXCR4, we also wondered how the co-application of MIF and MDLs might affect activation in the CXCR2-dependent yeast reporter system. Similar to the effect seen for CXCR4, joint application of MIF and either MDL1 or MDL2 resulted in a hyper-activation, indicating that synergistic activation involving the MIF and MDL mixtures also occurs for CXCR2 (Fig.3D).

We next used pharmacological probes to verify these surprising results. The MIF small molecule inhibitor ISO-1 (4,5-dihydro-3-(4-hydroxyphenyl)-5-isoxazoleacetic acid methyl ester) binds to the tautomerase pocket of MIF, thereby inhibiting its catalytic activity as well as its CD74-mediated induction of MAPK activation, p53-dependent apoptosis and cell proliferation (*43–45*). ISO-1 was previously also shown to partially block the MIF-CXCR4 reporter activation (*39*) and MDL1-induced monocyte chemotaxis (*33*), indicating that this inhibitor might likewise affect MDL1/CXCR4 effects. In the yeast-based CXCR4 reporter system, co-application of ISO-1 (100 µM) with MIF, MDL1 or MDL2 strongly reduced the activating capacity of these proteins (Fig. 3E). Notably, we also noticed a marked reduction of the synergistic effect triggered by the joint presence of MIF and MDL1 or MDL2 by ISO-1. The FDA-approved drug AMD3100 is a CXCR4 receptor antagonist that prevents the binding of CXCR4 ligands such as CXCL12, and partially inhibits MIF, thus constraining CXCR4 signaling (*25*). Using AMD3100 in the yeast reporter assay at a 10-fold molar excess over the concentration of the tested ligands, we observed significantly reduced CXCR4 activation by MIF and MDL2, both in single application and in combination of the two proteins. Interestingly, for MDL1 alone there was no inhibition by AMD3100, and only a mild reduction in signaling when co-applied with MIF (Fig. 3F). The CXCR2 antagonist SB225002 (*46*) (used at 20-fold molar excess over the ligands), reduced activation by MIF, MDL1 and MDL2 to similar degrees (Fig. 3G). Together, the results of these experiments showed that MDL1 and MDL2 are better agonists than MIF when used alone, and induce synergistic hyper-activation of CXCR2 and CXCR4 when used in combination with MIF, which can be largely blocked by MIF-specific small molecule inhibitors.

### MIF-MDL hetero-oligomer formation and its binding affinities

MIF can form various types of homo-oligomers but a monomer has never been reported (*24, 37, 47, 48*). This prompted us to investigate if MIF and MDLs can also form hetero-oligomers, which could be the basis of the observed remarkable synergistic effect on CXCR2 and CXCR4 receptor activation described above. Individual MIF and MDL1 proteins eluted as trimers in size exclusion chromatography (SEC), but a mixture of MIF and MDL1 showed formation of potential hexamers in addition to individual trimers (Fig. 4A, B). Elution volumes (Ve) and protein markers were used to obtain a standard curve (SFig. 8), which allowed estimations for the molecular masses of MIF (43.8 ± 0.7 kDa) and MDL1 (38.0 ± 0.3 kDa), as well as MDL2 (35.9 ± 0.6 kDa), when the proteins were applied individually (Table 1). These masses are well in line with the respective trimers. The estimated molecular masses obtained for SEC analysis of the MIF and MDL1 mixture were determined to be 38.5 ± 0.7 kDa and 82.5 ± 0.6 kDa (Table 1), values that are in good agreement with the molecular masses of a (homo-or heteromeric) trimer and a hetero-oligomeric hexamer, respectively. We noticed that only approximately one third of the MIF and MDL1 mixture formed a hetero-hexamer (Fig. 4B). To establish whether this proportion of the hetero-hexamer had any functional role, we used the molecule p425, a sulfonated azo compound proposed to support the formation of hexamers from MIF trimers (Fig. 4C). p425 was also shown to inhibit MIF tautomerase and CD74 activities (*47, 48*). We tested whether p425 affects hetero-hexamer formation between MIF and MDLs using the MST assays. In the presence of 100 µM of p425, the opposite effect was observed with p425, preventing binding of MIF to MDL1 (Fig. 4D), MDL2 and MDL3 (SFig. 7A, B). To investigate whether the trimer or hexamer contributed to the observed activation and synergism in the yeast-based assay, we used p425 (100 µM) in the CXCR4 signaling assay with application of the individual agonists MIF, MDL1, or MDL2 alone, or with co-application of MIF with MDL1 or MDL2.

**Figure 4.**
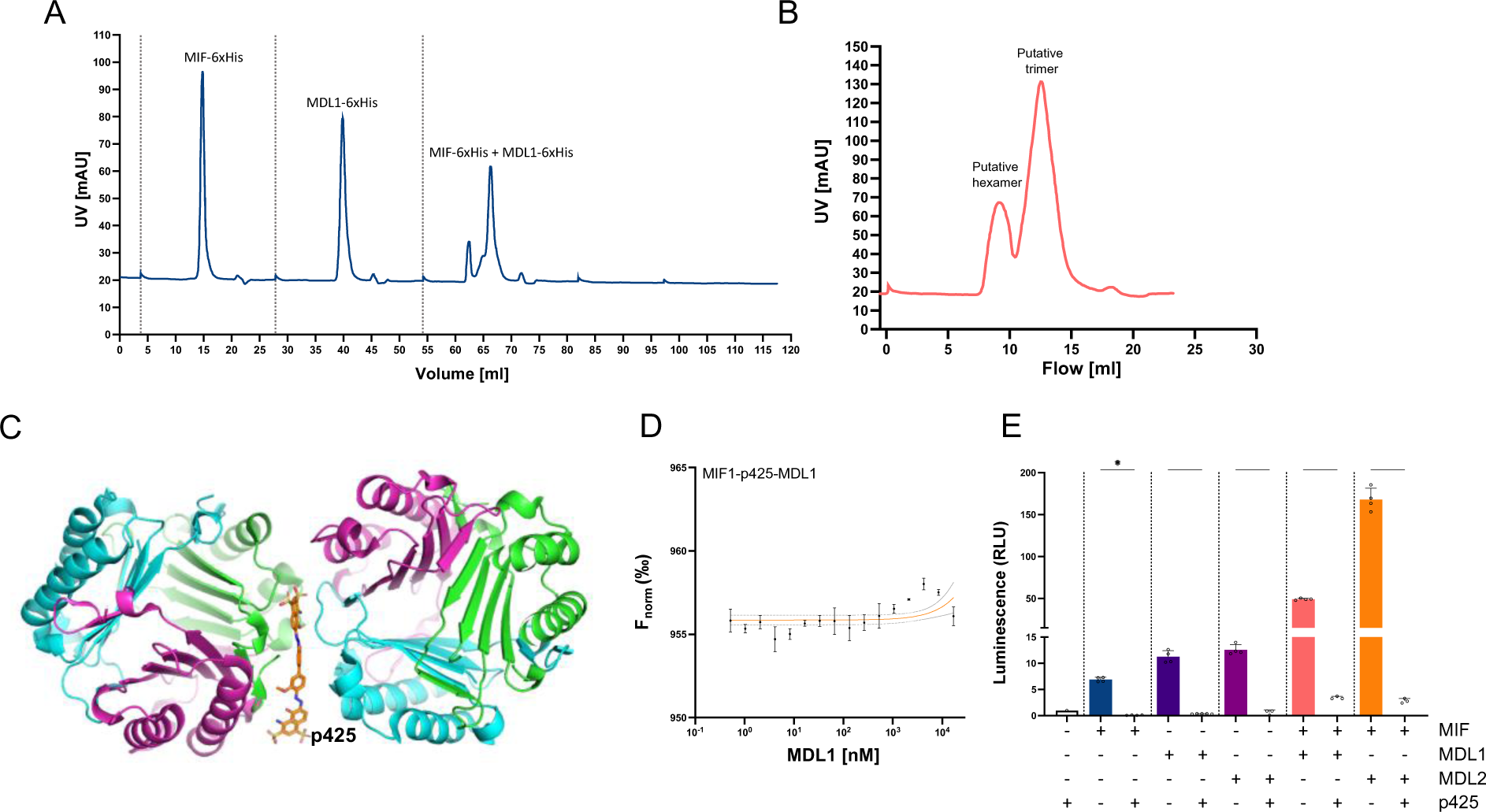
Direct protein-protein interaction via formation of higher-order oligomeric complexes between MIF and MDL1. (**A**) SEC chromatogram of individual MIF-6xHis and MDL1-6xHis and a mixture of MIF-6xHis each in 20 mM sodium phosphate, pH 7.4, at a constant flow rate of 0.5 mL/min. Depicted is the UV absorbance in mAU over the flow in mL. (**B**) SEC chromatogram showing a detailed view of the observed peak for a hexamer and trimer in the MIF and MDL1 mixture. (**C**) The crystal structure of p425 showing interactions between two trimers. (**D**) Direct protein–protein interaction studies between fluorescently labeled RED-NHS-MIF and MDL1 using microscale thermophoresis (MST). Inhibitor p425 was used at a 10-fold excess to MIF. For a constant MIF concentration of 100 nM, the difference in normalized fluorescence [‰] is plotted against increasing MDL1 concentrations for analysis of thermophoresis. Values shown represent means ± SD as obtained from at least three independent experiments. Further control experiments are shown in SFig. 7. (**E**) Effect of p425 on MIF/MDL-mediated chemokine receptor binding and signaling activation in the yeast-based CXCR4 reporter system. The plot shows luminescence (in RLU) due to *lacZ* reporter gene activation following the addition of MIF and MDL recombinant proteins, alone or in 1:1 combination, at a final total protein concentration of 20 µM, in the absence or presence of 100 µM p425. Values shown represent means ± SD as obtained from at least three independent experiments with RLUs of each experiment assessed in technical duplicates and normalized to untreated controls. Individual data points are indicated by white circles. Statistical analysis was performed using one-way ANOVA with multiple comparison (* p < 0.05, **** p < 0.0001).

**Table 1.**
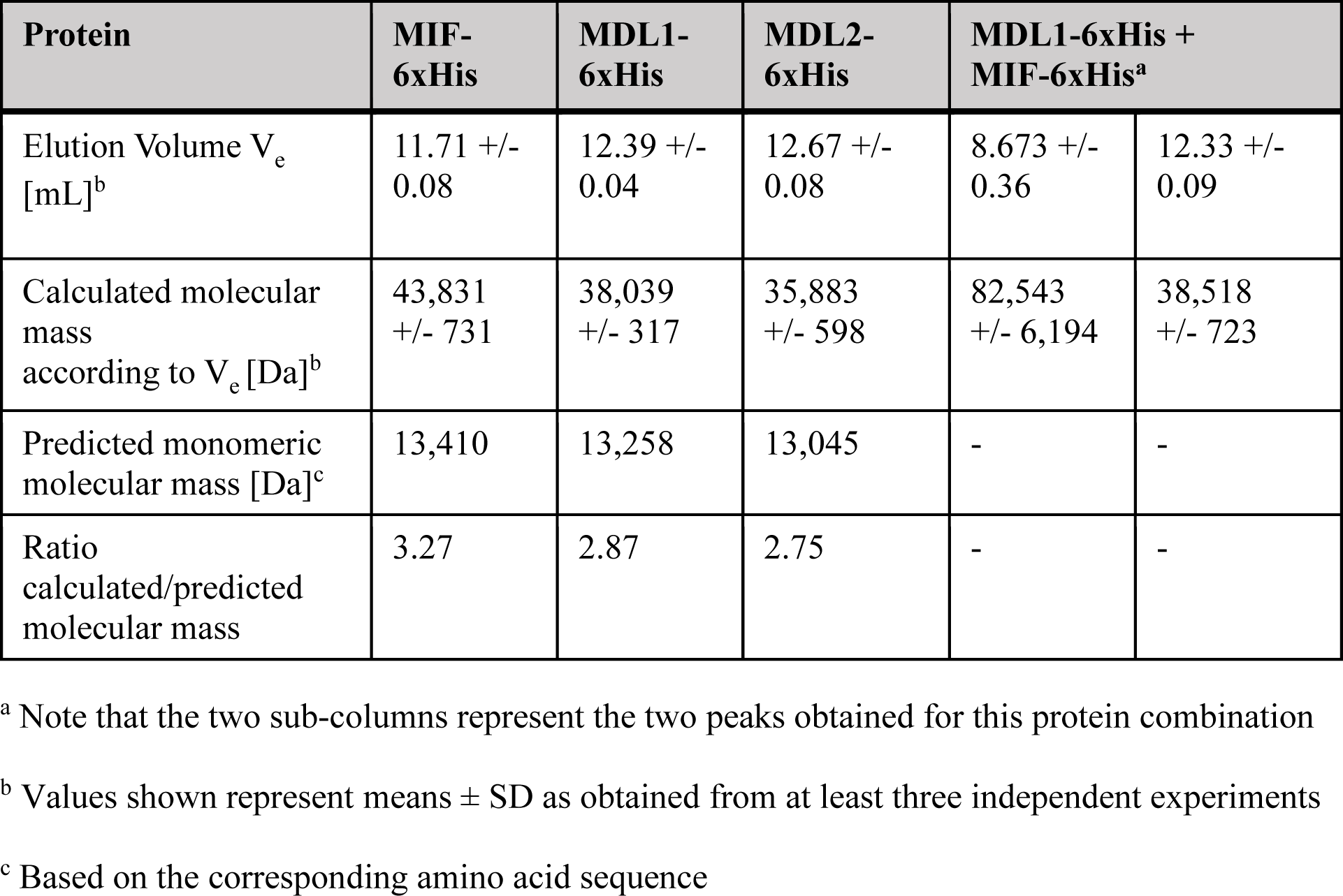
Chromatography statistics for SEC of recombinant MIF-6xHis and MDL-6xHis proteins.

Signaling activity was completely abolished in the case of the individual proteins, suggesting that activation of CXCR4 requires access to the tautomerase site, similar to the MIF small molecule inhibitor ISO-1 (Fig. 2 E) (*25*). Moreover, co-administration of p425 in the context of co-activation of MIF with MDL1 or MDL2 significantly reduced CXCR4 synergistic activity to basal levels (Fig. 4D). This result, together with the SEC and MST data, strongly suggest that the observed synergism is due oligomerization between a trimeric MIF and a trimeric MDL as the main molecular species responsible triggering CXCR4 signaling.

## DISCUSSION

In this study, we investigated structural, biochemical, and functional properties of MDLs, plant orthologs of the atypical human cytokine MIF. Analysis of the structural data obtained for all three MDLs showed an extraordinary degree of structural conservation at the enzymatic site, thereby confirming previous sequence-based *in silico* modelling (*31*). Despite the high degree of conservation at the tautomerase active site, a striking difference in enzymatic catalysis was observed in each subunit for all three MDLs, where Lys-98 has no stabilizing interaction with modelled HPP. There are other residues that are different in the MDLs, such as Phe-96 and Ile-96 for MDL2 and MDL3, respectively, but these are unlikely to have any effects on activity due to the large influence of Lys-98 (*33*). However, we cannot eliminate the possibility that binding of HPP to the MDL-1 or MDL-2 enzymatic site occurs in a non-productive manner for catalysis, as the MIF inhibitor ISO-1 was designed based on the MIF-HPP structure (*36*) and inhibits MDL-mediated activities.

We present evidence for direct protein-protein interaction and cooperative signaling of MIF with MDLs, tested in a variety of different systems including immunohistochemistry, yeast two-hybrid assays, binding experiments with MST, *in planta* experiments, CXCR2 and CXCR4 signaling experiments, and inhibition of signaling by pharmacological agents affecting MIF, chemokine receptors, and MIF-MDL oligomerization. Although plants do secrete proteins, there is no evidence that MDLs are exported outside the cell (*34*). Furthermore, there are no proteins resembling the MIF receptors for MDLs to activate these types of receptors in plants. Although the absence of known receptors does not dismiss activation of other receptors or intracellular proteins, evidence points to MDLs functioning as intracellular (or peroxisomal for MDL3) enzymes. To understand the role of MDLs in plant life, the respective physiological substrate needs to be identified for each MDL.

While the MST, yeast, and *in planta* assays do not present stoichiometric information, the SEC experiment suggests that a dimer of two different MIF/MDL homo-trimers is formed. This conclusion is supported by experiments using the inhibitor p425, which prevented or disrupted oligomerization, blocked MIF/MDL binding in the MST assay, and attenuated hetero-oligomer-mediated synergism in the CXCR4- or CXCR2-engineered strains of *S. cerevisiae*. Its mode of action involves intercalation between the interface region of two MIF trimers, thereby inhibiting MIF-mediated inflammatory responses (*47–49*). We would eliminate the possibility that there is a mixture of MIF and MDL within a trimer due to the dissociation rate of 7.7 × 10^-16^ M^2^ for MIF as determined by sedimentation velocity and equilibrium experiments (*50*). While the dissociation rates of the MDLs have not been measured, we assume, in analogy to MIF, they function as tight trimers and that the MDL1-MDL2 association observed in the two-hybrid assay is also based on an oligomer of homo-trimers, which might be an important information for determining the functional role of MDL1-MDL2 complexes in plants.

Given the results of the protein-protein interaction experiments, we probed whether there was signaling activity in a genetically modified strain of *S. cerevisiae* previously shown to express functionally active CXCR4 (*25, 39–42*). In addition to CXCR4, in this study CXCR2 was also used for analogous experiments. Both MDL1 and MDL2 induced signaling through CXCR2 and CXCR4. The MIF inhibitor ISO-1 inhibited CXCR4 signaling by both MDLs. Interestingly, antagonists of CXCR2 (SB-225002) and CXCR4 (AMD3100) have different effects on MDL1- and MDL2-mediated signaling. While SB-225002 and AMD3100 inhibited activation of CXCR2 and CXCR4 by MDL2, respectively, there was no effect of AMD3100 on CXCR4 signaling by MDL1, suggesting an allosteric mechanism of MDL1 activation that bypasses the CXCR4 trans-membrane cavity that is necessary for orthosteric activation (*51*). This finding thus also illustrates the plasticity of CXCR4 activation. What is surprising is the synergy that occurs when MDL1 or MDL2 together with MIF co-stimulates either CXCR2 or CXCR4. In this case, the inhibitors (ISO-1, SB-225002, and AMD3100) decreased activation. To assess how these two MIF-MDL complexes synergize, the inhibitor p425, which dramatically reduced synergism to almost basal levels, provided initial insight. To gain greater mechanistic insight as to whether MIF and MDL1 and MDL2 associate or act independently to achieve signaling synergy, we used SEC to show that MIF and MDLs combine to form a putative trimer-trimer (hexameric) complex.

We realize the cross-kingdom interactions between human MIF and plant (A. thaliana) MIF homologues (termed MDLs) are unexpected, but the *A. thaliana* MDLs share high identity to many other plant MDLs (>90%) which are nutrients or spices for humans such as cabbage, broccoli, cauliflower, kale, Brussels sprouts, collard greens, radish, and wasabi among many others, and are anticipated to have the same effects with human MIF and its receptors. The interactions among MDLs, MIF, and signaling through human receptors is a novel finding and may be representative of other molecules within nutrients during digestion. A close alternative is well known that plant extracts from “medicinal plants” that can have immunomodulatory activities on mammals, with the mechanistic basis remaining poorly understood.

Immunomodulatory effects of plant extracts may relate to the presence of phytochemicals, for example plant-derived secondary metabolites such as flavonoids, polysaccharides, lactones, alkaloids, diterpenoids or glycosides (*52, 53*). Plant-derived peptides (*54*) and proteins (*55*) have been likewise proposed to affect mammalian immune status. Although *A. thaliana* is neither a “medicinal plant” nor used as nutrient in humans, the highly sequence-related MDL orthologs are omnipresent in other species of the plant kingdom (*31*). We propose that mammalian MIF activity could be modulated by direct contact with plant-derived MDL proteins such as following inhalation of plant particles or upon food ingestion. Additional *in vitro* and *in vivo* studies are necessary to investigate further the concept of modulation of the human immune system by plant MDL proteins, particularly in the context of MIF-driven diseases such as acute or chronic inflammation. Such studies involving MIF and MDLs are needed to broaden our understanding of these proteins in cross-kingdom interactions.

## MATERIALS and METHODS

### Expression and purification of recombinant proteins

Clones of MIF and the three *A. thaliana* MIF ortholog genes, *MDL1*, *MDL2*, and *MDL3* in pET21a were previously generated (*56*) and used in this work. Briefly, classical cloning strategies were applied and all genes were N-terminally fused in-frame to a hexahistidine tag included in the pET21a vector using the restriction endonucleases *Nde* I and *Xho* I. Plasmids were transformed into competent *Escherichia coli* Rosetta^TM^ (DE3) cells to express the pET21-derived genes and to yield MIF–6xHis, MDL1–6xHis, MDL2– 6xHis, and MDL3–6xHis fusion proteins. Culturing and induction of protein expression following application of the inductor isopropyl-β-D-thiogalactopyranoside (IPTG; Sigma, Deisenhofen, Germany)) was performed as previously described (*33*).

To release intracellular protein, a high-pressure cell homogenizer (French press, Avestin EmulsiFlex C5 by Avestin Europe GmbH, Mannheim, Germany) was used to lyse cells at approximately 75 MPa. Homogenization as well as all following purification steps were carried out on ice and under constant cooling. For homogenization, fresh or frozen bacterial pellets gently thawed on ice were resuspended in 1 mL ice-cold immobilized metal affinity chromatography (IMAC) binding buffer (20 mM sodium phosphate, 0.5 M NaCl, 20 mM imidazole, pH 7.2). Lysates were then centrifuged at 18.000 × *g* for 30 min at 4 °C to remove cell debris. The protein-containing supernatants were collected and filtered prior to usage in fast protein liquid chromatography (FPLC).

For purification, IMAC and subsequent SEC were performed on an FPLC system (ÄKTA Pure, GE Healthcare/Cytiva, Freiburg, Germany). Nickel-loaded IMAC columns (HisTrap, GE Healthcare/Cytiva) equilibrated with at least 5 column volumes of IMAC binding buffer were loaded with protein lysates under a flow rate of 1 mL/min binding buffer. His-tagged protein was then eluted by a gradient over 30 min, flow rate 0.5 mL/min from 0% to 100% IMAC elution buffer (20 mM sodium phosphate, 0.5 M NaCl, 0.5 M imidazole, pH 7.2). During elution, samples were collected in fractions of 0.5 mL and protein content monitored via an UV-detector at 280 nm. Protein-containing fractions were combined and purified further via SEC on a Superdex 75 10/300 GL column (GE Healthcare/Cytiva) using 20 mM sodium phosphate buffer, pH 7.2, a buffer condition previously reported to preserve MIF bioactivity (*56*). Protein-containing and imidazole-free fractions were collected and sterile-filtered over a 0.2 µm filter prior to further use. Protein purity was assessed by sodium dodecylsulfate-polyacrylamide gel electrophoresis (SDS-PAGE) with Coomassie and silver staining as well as anti-6xHis immunoblot (see below). Endotoxin content was measured photometrically in final, sterile-filtered protein solution using the Pierce LAL Chromogenic Endotoxin Quantitation Kit (Thermo Fisher Scientific, Dreieich, Germany) essentially following the manufacturer’s instruction. Purified protein was stored at 4 °C and used within a maximum of 4 weeks.

### Crystallization and structure determination

For crystallization, buffers of all MDL proteins were exchanged for 20 mM HEPES, 250 mM NaCl, pH 7.5, immediately after purification and the protein was then concentrated (MDL1: 12.3 mg/mL, MDL2: 11.1 mg/mL, MDL3: 9.8 mg/mL). For all MDL proteins, crystallization experiments were carried out using sitting drop/vapor diffusion crystallization in a 200 nL + 200 nL format using a Phoenix crystallization robot in MRC-2 crystallization plates. Individual crystallization conditions for MDL1 consisted of 50 mM Tris, pH 8.0, 0.2 M calcium acetate, and 26% PEG 8000. For MDL2, the crystallization conditions were 50 mM Tris, pH 8.0, and 2.25 M ammonium sulfate; and for MDL3 they were 50 mM MES, pH 6.0, 4% MPD, 0.2 M ammonium acetate, and 30% PEG3350. All crystals were grown at 292.15 K. For cryoprotection, individual crystals were transferred to a new drop containing the mother liquor enriched with 30% ethylene glycol for MDL1, 30% glycerol for MDL2, and 30% ethylene glycol for MDL3, and flash-frozen in liquid nitrogen. X-ray data were collected at 100 K at the Paul Scherrer Institute (PSI) synchrotron using a Dectris Eiger2 16M Detector (wavelength=0.9999 Å).

Diffraction data reduction was done using the XDS program (*57*). The observed reflections were scaled and merged using Aimless (*58*) provided in the CCP4 software. The crystal structures were solved by the phaser molecular replacement method using Phenix-GUI (*59*). The MIF monomer structure (PDB 7KQX) was used as a model for structure determination. The structure solution yielded a trimer in the asymmetric unit for MDL1 and MDL2 and a monomer for MDL3. The refinement of the structures was performed using the module Phenix.refine (*60*) of the PHENIX package. Cycles of refinement and model building were performed using Phenix.refine and Coot (*61*). The stereochemistry of these crystal structures was assessed using MOLPROBITY (*62*). Individual refinement statistics for each protein are listed in Supplementary Table 1. All structural data were collected at 1.56 Å, 1.40 Å, and 2.0 Å for MDL1, MDL2, and MDL3, respectively, deposited in the RCSB (PDBs 8DQA, 8AP3, 8DQ6 for MDL1, MDL2, MDL3, respectively). The crystal structures were compared to each other and to the previously published MIF structure (PDB 3DJH; (63)) with PyMOL and Chimera software (64) used for visualization and analysis.

### Generation of monoclonal anti-MDL1 antibodies

Lou/c rats were immunized with 60 µg purified full-length MDL1-6xHis protein, 5 nmol CpG (TIB MOLBIOL, Berlin, Germany), and an equal volume of Incomplete Freund’s adjuvant (IFA; Sigma, St. Louis, USA). Hybridoma supernatants were generated and screened as described previously (*34*). Selected supernatants were validated by slot blot immunoassay on recombinant purified human MIF (MIF), human MIF-2/D-DT, and the three MDL proteins for specificity and sensitivity (SFig. 5). Hybridoma cells from clone ATM1 21G9 (IgG2b/ƙ) were subcloned twice by limiting dilution to obtain a stable monoclonal antibody-producing cell line.

### SDS-PAGE and immunoblot analysis of recombinant MDL-6xHis proteins

Following purification, protein purity was assessed via SDS-PAGE and Coomassie staining or silver staining and anti-6xHis immunoblotting. Electrophoresis was performed in 15% acrylamide gels under reducing conditions as described before (*33*). For immunoblot analysis, electrophoresed proteins were transferred to nitrocellulose membranes using Tris-glycine transfer buffer (Thermo Fisher Scientific), followed by blocking (1% BSA) and staining in TBST (Tris-buffered saline, 150 mM NaCl, 20 mM Tris, 0.01% Tween-20, pH 7.3) supplemented with 1% bovine serum albumin (BSA) (Sigma-Aldrich). Hexahistidine-tagged proteins were then detected using mouse anti-6xHis tag antibody (Ma1-135, Invitrogen, Karlsruhe, Germany) as primary antibody and signals revealed by horseradish peroxidase (HRP)-conjugated goat anti-mouse IgG (ab6789, Abcam, Cambridge, UK). Imaging was performed upon addition of SuperSignal^TM^ West Dura Extended Duration Substrate (Thermo Fisher Scientific) on an Odyssey Fc Imaging System using ImageStudioTM software (LICOR Biosciences, Bad Homburg, Germany).

### Co-immunoprecipitation

Prior to immunoprecipitation, MIF-6xHis was biotinylated using EZ-Link™ Sulfo-NHS-LC-Biotinylation Kit (Thermo Fisher Scientific), performed essentially as per manufacturer’s instructions. For biotinylation, 1 mg recombinant MIF-6xHis at a concentration of 1 mg/mL in 20 mM sodium phosphate buffer, pH 7.2, was used. Co-immunoprecipitation experiments were then carried out using Dynabeads^TM^ M-280 streptavidin (Thermo Fisher Scientific). To allow time for interaction, recombinant MIF-6xHis-Biotin and MDL1-6xHis were mixed at an equimolar ratio and incubated overnight at 4 °C. The next day, beads were resuspended thoroughly and 50 µL beads were washed with 1 mL washing buffer (PBS, 0.01% BSA, pH 7.4). Following magnetic isolation and resuspension in 50 µL washing buffer, beads were added to the protein mixture and incubated with slight agitation for 30 min at room temperature. Thereafter, beads and all protein bound to them were magnetically isolated for 3 min, with the supernatant then removed. Beads were washed 3 times with 1 mL washing buffer, resuspended in 50 µL denaturing SDS-PAGE sample buffer and boiled for 5 min at 95 °C prior to analysis by SDS-PAGE.

Immunoblotting was performed as described above and MIF-6xHis-biotin revealed using streptavidin-HRP-conjugate (Merck, Darmstadt, Germany), while MDL1-6xHis was revealed using a custom-made monoclonal primary antibody (1:10 dilution), specifically designed to distinguish between MIF and MDL1 (see above), and HRP-coupled subclass-specific monoclonal anti-rat secondary antibody (RG7/11.1; ATCC).

### Yeast two-hybrid binding assay

For yeast two-hybrid assay, Gateway® cloning-compatible vectors pDEST32 and pDEST22 (Invitrogen ProQuest yeast two-hybrid System) were used, which enable N-terminal fusions of bait and prey proteins with the Gal4 activation- and DNA-binding domains, respectively. *MIF* and *MDL* coding sequences were mobilized from pDONR207 entry clones via Gateway^®^ recombination into pDEST32 and pDEST22. The resulting plasmids were transformed into *S. cerevisiae* strain PJ69-4A (*65*). Yeast transformants were dropped on appropriate synthetic complete media lacking selective amino acids for growth control and detecting putative interactions. For drop tests, yeast cultures were grown overnight, washed with sterile water, adjusted to an OD600 of 1, ten-fold dilution series established, and 4 µl per strain and dilution dropped onto the corresponding medium. Bait and prey protein expression was validated by immunoblot analysis using the GAL4 (DBD) (SC-510) and GAL4 (AD) (SC-1663) monoclonal antibodies (Santa Cruz Biotechnology, Dallas, TX, USA).

### Luciferase complementation imaging (LCI) assays

For LCI assays, Gateway^®^ cloning-compatible vectors pAMPAT-nLUC-GWY and pAMPAT-cLUC-GWY (*34*) were used, which enable N-terminal fusions of bait and prey proteins with the N- and C-terminal segments of firefly luciferase, respectively. *MIF* and *MDL* coding sequences were mobilized from pDONR207 entry clones via Gateway^®^ recombination into pAMPAT-nLUC-GWY and pAMPAT-cLUC-GWY. The resulting plasmids were transformed into *A. tumefaciens* strain GVG3101 (pMP90RK). Bacterial cultures were grown overnight, resuspended in infiltration media (10 mM MES, pH 5.6, 10 mM MgCl2, 200 µM acetosyringone) to an OD600 of 0.5 and incubated at room temperature for 2 h. For co-infiltration, equal volumes of each *A. tumefaciens* transformant were mixed and infiltrated with a needleless syringe from the abaxial side into fully expanded leaves of four- to six-week-old *N. benthamiana* plants. The leaves were sprayed with 1 mM D-luciferin (PerkinElmer) solved in water supplemented with 0.01% (v/v) Tween-20 at three days after infiltration. Leaves were kept in the dark for 10 min before luminescence was detected with a ChemiDoc™ XRS+ imagine system (Bio-Rad, Feldkirchen, Germany). Luminscence intensities/mm^2^ infiltrated leaf area of different combinations were evaluated using the Image Lab software (BioRad, version 6.1). For each combination of interaction partners, three independent experiments consisting of two different plants and two leaves per plant were evaluated. *Agrobacterium*-mediated transient expression of LCI constructs in *N. benthamiana* was validated by immunoblot analysis using a polyclonal anti-luciferase primary antibody (Merck; used in 1:1000 dilution).

### Chemokine receptor signaling assay in yeast

For receptor signaling experiments, we used the functional CXCR4- or CXCR2-expressing transformants of *S. cerevisiae* strain CY12946 that has been previously describe (*25, 39–41*). Briefly, the endogenous yeast pheromone receptor was replaced by human CXCR4 or CXCR2, respectively, with the activated human chemokine receptor being functionally linked to the downstream MAPK-type signaling pathway, ultimately resulting in expression of the *lacZ/β-gal* reporter gene upon receptor binding. The β-galactosidase enzymatic activity (assessed photometrically) was therefore used as a surrogate parameter for chemokine receptor activation.

Yeast cells were grown in a 24-well plate until reaching an OD600 of 0.3-0.8 and then incubated with the respective protein samples (MDL1-6xHis, MDL2-6xHis, MDL3-6xHis) or the known agonist MIF-6xHis, either individually or as combinations, either with or without inhibitors added. A concentration of 10-20 µM protein has previously been shown to create stable responses and was used as a reference point for the inhibitor studies. It must be noted that, due to the barrier function of the yeast cell wall, high doses are needed for stable receptor activation.

Activation of chemokine receptors was detected by measuring ß-galactosidase activity using the commercially available Beta-Glo assay system (Promega Corp, Madison, WI, USA) and luminescence signal was recorded on a multimodal plate reader (Enspire 2300, PerkinElmer Life Sciences, Rodgau, Germany). The kit was used as per manufacturer’s instructions and is based on coupling ß-galactosidase enzymatic activity to a luciferase reaction. After mixing assay buffer and assay substrate in a 1:1 ratio, a volume of this mixture equal to the media volume was added to each well. After mixing and incubation at room temperature for 30 min, luminescence of each sample was measured.

### Microscale thermophoresis (MST)

All MST experiments were performed on a Monolith NT.115 instrument with green/red filters (NanoTemper Technologies, Munich, Germany). MST and LED power were set at 40% and 60%, respectively, for MDL1 and MDL2, or 40% and 90% for MDL3 measurements to obtain stable fluorescent signals around 1000 fluorescent counts. All measurements were performed at 37 °C with MST traces tracked for 40 s (laser-off: 5 s, laser-on: 30 s; laser-off: 5 s). A stock solution of 200 nM RED-NHS-MIF-6xHis was prepared in 20 mM sodium phosphate buffer, pH 7.4, containing 0.2% BSA, according to the manufacturer’s protocol.

For titration of each plant ortholog, each protein sub-stock solution was prepared by serial 1:1 dilutions, starting from a 20 µM stock solution in 20 mM sodium phosphate buffer, pH 7.4, 0.1 % BSA. RED-NHS-MIF-6xHis and each MDL sub-stock was mixed at a 1:1 ratio resulting in a final MIF concentration of 100 nM and incubated for 10 min in the dark at room temperature. Since initial screening had shown slight sticking of protein to standard capillary walls, premium coated capillaries were used throughout. Incubated mixtures were loaded into capillaries and MST measurements started immediately. Obtained MST traces were analyzed at an MST-on time of 1.5 s using the MO.Affinity Analysis version 2.2.4 (NanoTemper Technologies) for each of the three potential interaction pairs. Apparent KD values were calculated using the same software. Visualization was done using Prism GraphPad (Version 9.4.1) assuming a 1 on 1 binding model with sigmoidal curve fitting models for each set up.

### Size Exclusion Chromatography

SEC experiments were performed on an FPLC system (ÄKTA Pure, GE Healthcare/Cytiva, Freiburg, Germany) with a Superdex 75 10/300 GL column (GE Healthcare/Cytiva) using 20 mM sodium phosphate buffer, pH 7.4, and a constant flow of 0.5 mL/min. Protein were used for SEC 24 hours after purification, either individually or in a 1:1 mixture of MIF-6xHis and MDL1-6xHis, incubated at 4 °C overnight. Proteins were loaded individually, one after another, and peaks observed by UV absorbance (280nm) in mAU over the elution volume in mL. Unicorn 7.0 software (GE Healthcare/Cytiva, Freiburg, Germany) was used to analyze chromatograms for individual elution volumes. Experiments were performed in triplicates.

For the described SEC setup, a stand curve and standard equation was generated using the GE gel filtration calibration kit, LMW, as per manufacturer’s instructions (GE Healthcare/Cytiva, Freiburg, Germany). From observed elution volumes and known molecular mass of sample proteins, a standard curve and standard equation were calculated and visualized using Prism GraphPad (Version 9.4.1).

### Statistics

Unless otherwise indicated, statistical analyses were performed using one-way analysis of variance (ANOVA) followed by *post hoc* comparison with the Bonferroni test using GraphPad Prism 9 (GraphPad Prism Software Inc., San Diego, CA) with multiple comparisons. Data are presented as means ± SD. Considered as significant: p < 0.05. Asterisks indicate statistically significant differences as follows: *, p < 0.05; **, p < 0.01; ***, p < 0.005; ****, p < 0.001.

## Acknowledgements

We thank Elena Conti for making arrangements at the Max-Planck-Institut für Biochemie for X-ray data collection and processing, and Simona Gerra for valuable technical help with the expression of recombinant MIF and MDL proteins.

## Funding

This work was supported by the Deutsche Forschungsgemeinschaft (DFG, German Research Foundation)-Agence Nationale Recherche (ANR) co-funded project “X-KINGDOM-MIF - Cross-kingdom analysis of macrophage migration inhibitory factor (MIF) functions”. Respective DFG grants are BE 1977/10-1 to J.B., PA 861/15-1 to R.P. The monoclonal antibody facility and the Division of Vascular Biology of LMU are co-funded by the DFG under Germanýs Excellence Strategy within the framework of the Munich Cluster for Systems Neurology (EXC 2145 SyNergy; ID 390857198). L.S. acknowledges receipt of a fellowship from the Studienstiftung des Deutschen Volkes.

## Author contributions

Conceptualization: EL, JB and RP

Methodology: LS, RM, FL, JB, PB, DS, MB, SL, BS, and AF

Data Analysis: LS, RM, JB, and Figures: LS and RM

Provided critical materials: DS, MB, and RF Supervision: RF, RP, JB, and EL

Writing – original draft: all authors

Writing – review and editing: LS, EL, JB and RP

### Competing interests

JB is a coinventor on patent applications related to anti-inflammatory strategies to target MIF; all other authors declare they have no competing interests.

### Data and materials availability

The PDB atomic coordinates for MDL1, MDL2, and MDL3 are accessible from the RCSB from files 8DQA, 8AP3, and 8DQ6, respectively.

## Supplemental Information

**Supplemental Table 1.** Data collection and refinement statistics.

|  | **MDL1 (8DQA)** | **MDL2 (8AP3)** | **MDL3 (8DQ6)** |
| --- | --- | --- | --- |
| **Wavelength (Å)** | 0.9999 | 0.9999 | 0.9999 |
| **Resolution range (Å)** | 43.53-1.56 (1.616-1.56) | 37.32 -1.40 (1.45-1.40) | 44.75 -2.0 (2.072- 2.00) |
| **Space group** | P 65 | P 21 21 21 | I 2 3 |
| **Unit cell dimensions (Å)** | a, b=66.737,c=132.32<br>$\alpha=\beta=90^\circ$ $\gamma=120^\circ$ | a=46.218 b=79.243 c=84.6<br>$\alpha=\beta=\gamma=90^\circ$ | a, b, c=89.499<br>$\alpha=\beta=\gamma=90^\circ$ |
| **Total reflections** | 931420 (75211) | 783989 (72179) | 105554 (9776) |
| **Unique reflections** | 46283 (4257) | 61915 (6095) | 8229 (804) |
| **Multiplicity** | 20.1 (17.7) | 12.7 (11.8) | 12.8 (12.2) |
| **Completeness (%)** | 97.54 (90.40) | 99.96 (100.00) | 99.96 (100.00) |
| **Mean I/sigma(I)** | 31.45 (2.24) | 31.44 (3.01) | 35.38 (4.35) |
| **Wilson B-factor** | 26.64 | 19.22 | 40.22 |
| **R-merge** | 0.05281 (0.99) | 0.04017 (0.6908) | 0.04159 (0.5394) |
| **R-meas** | 0.05418 (1.019) | 0.04189 (0.7219) | 0.04334 (0.5634) |
| **R-pim** | 0.012 (0.236) | 0.01172 (0.2084) | 0.01205 (0.1612) |
| **CC1/2** | 1 (0.817) | 1 (0.907) | 1 (0.923) |
| **CC\*** | 1 (0.948) | 1 (0.975) | 1 (0.98) |
| **Reflections used in refinement** | 46163 (4256) | 61898 (6095) | 8227 (804) |
| **Reflections used for R-free** | 2308 (213) | 3173 (272) | 375 (37) |
| **R-work** | 0.1830 (0.2623) | 0.1847 (0.2678) | 0.2011 (0.2910) |
| **R-free** | 0.2083 (0.2874) | 0.2080 (0.3109) | 0.2324 (0.2579) |
| **CC(work)** | 0.967 (0.842) | 0.960 (0.864) | 0.957 (0.774) |
| **CC(free)** | 0.957 (0.815) | 0.949 (0.852) | 0.906 (0.927) |
| **Number of non-hydrogen atoms** | 2437 | 2823 | 784 |
| **macromolecules** | 2297 | 2602 | 752 |
| **solvent** | 134 | 221 | 32 |
| **Protein residues** | 309 | 351 | 100 |
| **RMS(bonds)** | 0.006 | 0.009 | 0.002 |
| **RMS (angles)** | 0.84 | 1.09 | 0.49 |
| **Ramachandran favored (%)** | 100.00 | 97.97 | 97.96 |
| **Ramachandran allowed (%)** | 0.00 | 2.03 | 2.04 |
| **Ramachandran outliers (%)** | 0.00 | 0.00 | 0.00 |
| **Rotamer outliers (%)** | 0.40 | 0.00 | 0.00 |
| **Clashscore** | 2.60 | 1.15 | 1.33 |
| **Average B-factor (Å<sup>2</sup>)** | 31.29 | 21.14 | 44.98 |
| **macromolecules** | 30.68 | 20.25 | 44.93 |
| **solvent** | 40.57 | 31.51 | 46.15 |

Statistics for the highest-resolution shell are shown in parentheses.

## SUPPLEMENTAL FIGURES

**Supplemental Figure 1.**
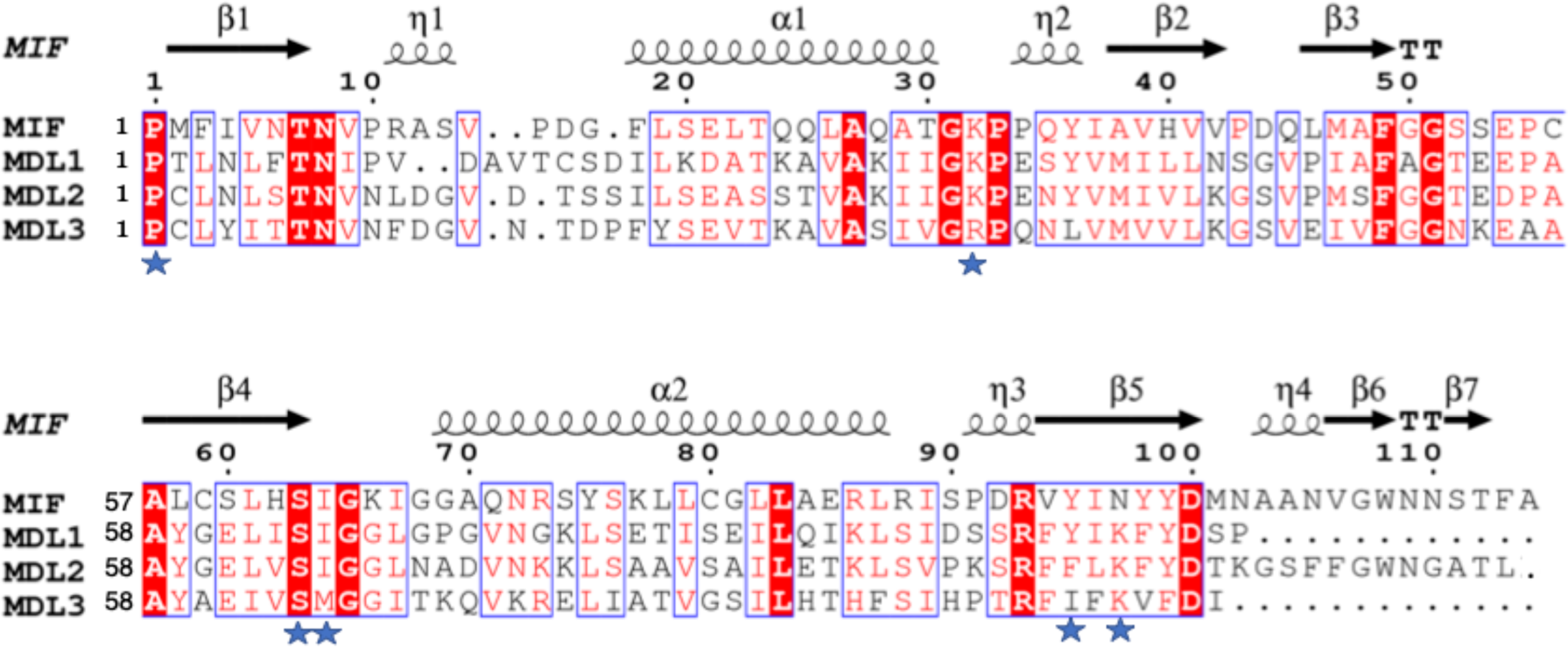
Structure-based sequence alignment of MDLs (accession numbers for MDL1, MDL2, and MDL3 are NP_001190557.1, NP_195785.1, and NP_566955.1, respectively) and human MIF. The ESpript application (1) was used to align the structures. The red boxes with a white character represent the invariant residues among the three MDLs and MIF, indicating evolutionary conservation between *A. thaliana* MDLs and human MIF. Similar residues or regions are surrounded by a blue box. The secondary structure elements are presented on top of the sequences, with the 310-helix represented by the η symbol, helices with squiggles, β-strands with arrows, and β-turns with TT letters. Blue stars below the aligned sequences indicate the position of MIF residues in the tautomerase catalytic site. Only residues with observed electron density in the structures are aligned; the last 12 and 9 residues of MDL1 and MDL3, respectively, are not visible. *Note regarding the numbering of the amino acids for MIF and MDLs:* (a) in some MIF studies, the first amino is referred as Met-1, representing the initiating methionine which is then post-translationally cleaved by methionine aminopeptidase, but in this and some other studies the first residue is numbered as Pro-1, referring to the first amino acid in the mature protein sequence; (b) after residue 17 for all four proteins, there is one extra amino acid in a loop for all three MDLs relative to MIF resulting in residue numbers for MDLs that are greater than MIF.

**Supplemental Figure 2.**
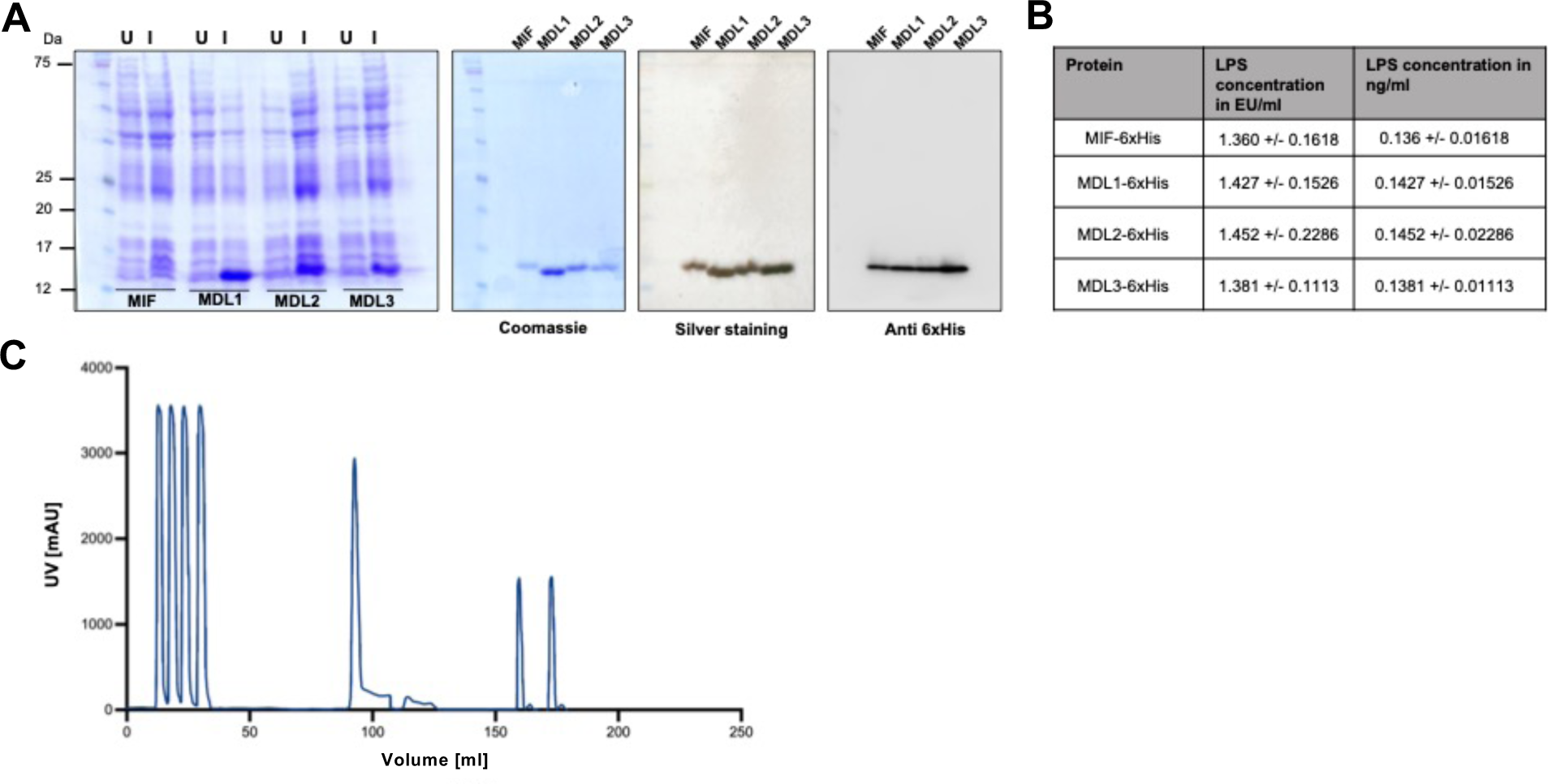
Expression und purification of recombinant MIF and MDL proteins. (**A**) Electrophoretic analysis of protein lysates before (undinduced [U]) and after induction [I] with isopropyl-β-D-thiogalactopyranoside (IPTG) (Coomassie staining). In addition, purified proteins are shown using Coomassie staining, silver staining, and anti-6xHis immunoblot. (**B**) Quantification of lipopolysaccharide (LPS) contamination in purified recombinant proteins using a chromogenic endotoxin detection assay. (**C**) Chromatogram of immobilized metal affinity chromatography (IMAC) and subsequent size exclusion chromatography (SEC) purification shown for MDL1 as an example. Injections of the bacterial lysate (up to 50 mL) are followed by elution of the His-tagged protein by an imidazole gradient (around 100 mL). For further purification and buffer exchange, this step is followed by two runs of SEC (from 150 mL onward).

**Supplemental Figure 3.**
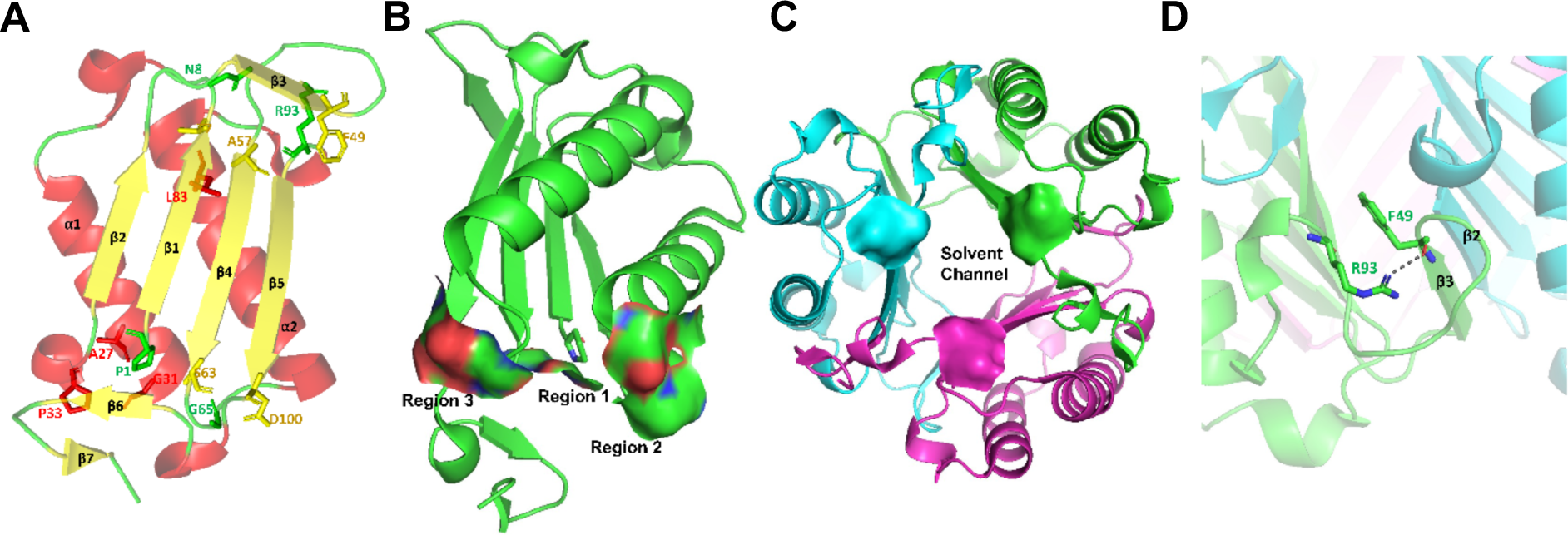
Structural views and regions of the invariant residues in MIF and MDL proteins. (**A**) All 14 invariant residues among the three MDLs and MIF are shown in a MIF monomer. See SFig. 1 for further reference. (**B**) Surface area of regions 1, 2, and 3. Region 1 contains the Pro-1 and Ser-3 of the tautomerase enzymatic site. Region 2 consists of Ala-27, Gly-31, Pro-33, Gly-65, Ser-63, and Asp-100. For Ser-63, the backbone atoms are in region 2, while the side chain is part of region 1. (**C**) The MIF trimer shows the solvent channel (water molecules not shown) along the 3-fold axis of the trimeric structure surrounded by three surface areas (shown as a smooth surface) of Asp-100 side chains from each subunit at one end of the channel and consist of region 3 and shown in different colors. (**D**) In region 4, a hydrogen between the side chain of Arg-93 and the backbone of Phe-49 stabilizes the β-strand important for subunit-subunit interactions (cartoons in blue and green represent two different subunits).

**Supplemental Figure 4.**
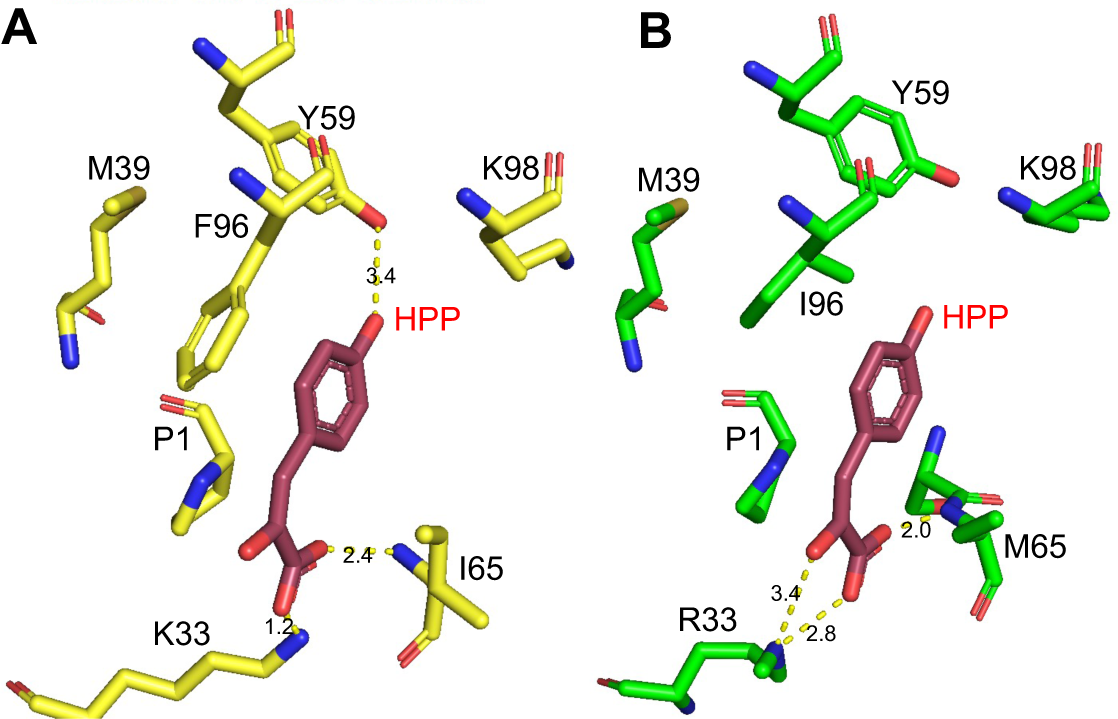
Interactions of MDL2 and MDL3 enzymatic site residues interacting with a modelled HPP. (**A**) MDL2 and (**B**) MDL3 catalytic residues in yellow and green, respectively, were superimposed on the MIF-HPP complex to examine the interactions between ligand and protein. Potential H-bonds are shown between the MDLs and HPP represented as yellow lines. Similar to Fig. 1C, in which the interaction between MDL1 and HPP is shown.

**Supplemental Figure 5.**
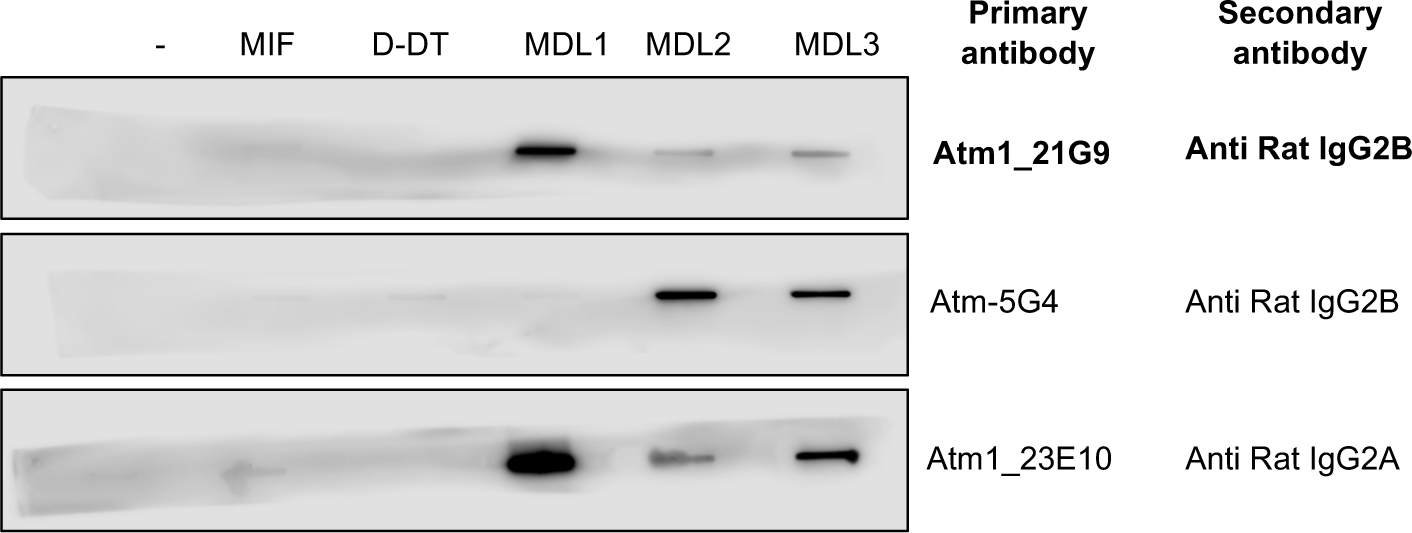
Slot blot screening for sensitivity and specificity of custom-made monoclonal antibodies against MDL1 and MDL2. Slot blot screening of custom-made monoclonal antibodies were examined against MDL1 and MDL2. Promising monoclonal antibodies directed against MDL1 and MDL2 were probed against recombinant purified human MIF (MIF), human MIF-2/D-DT (D-DT), and the three MDL proteins as indicated. The far left-hand lane was a negative control without loaded protein. HRP-coupled immunoglobulin subclass-specific secondary antibodies were used as indicated for detection. Antibody clones directed against MDL2 were previously established (2). Here, clone Atm-5G4 for MDL2 was used and is shown to recognize MDL3 in addition to MDL2. Two promising antibodies directed against MDL1 (Atm1_21G9 and Atm1_23E10) were probed. While both candidate antibodies were able to reasonably distinguish between MIF or D-DT and the MDLs, only Atm_21G9 showed a high degree of specificity for MDL1. Screening was performed with the primary hybridoma supernatant (shown here) and then validated with the established clone, which was used for subsequent experiments.

**Supplemental Figure 6.**
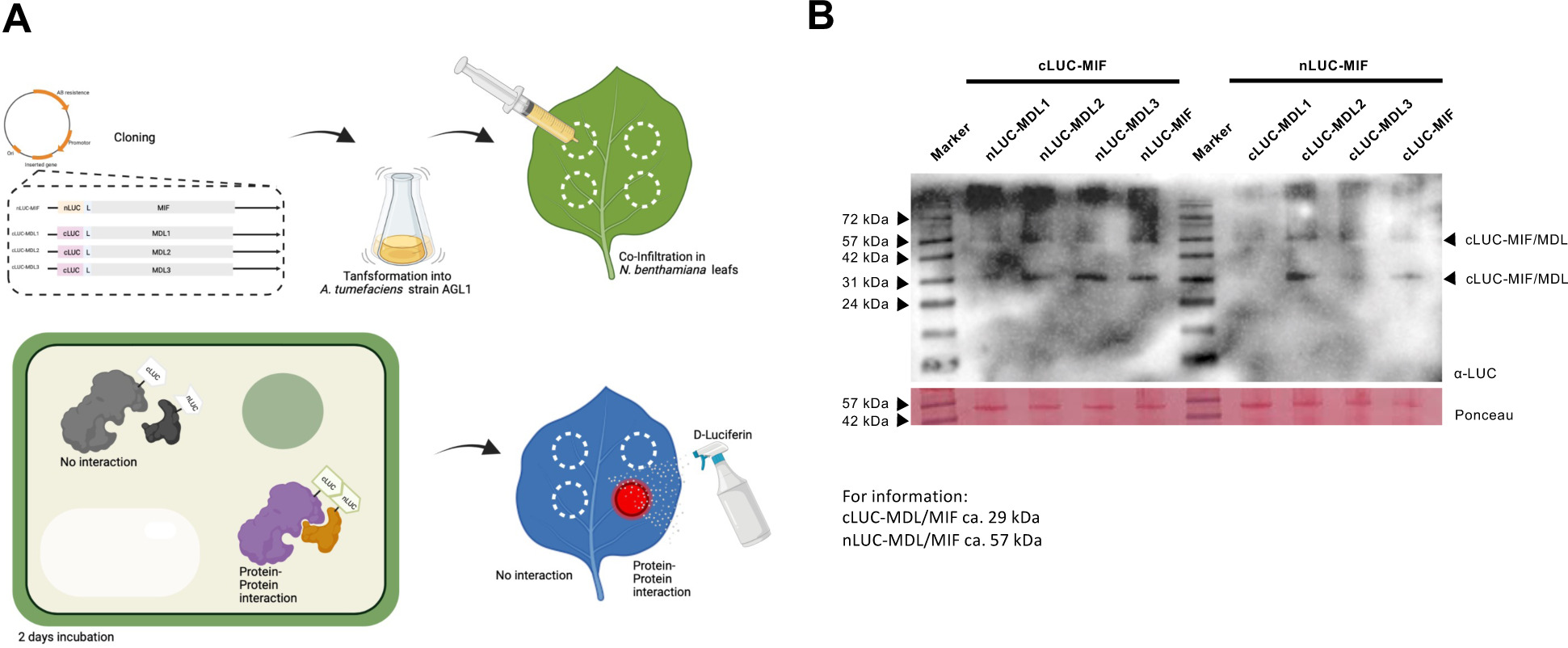
Luciferase complementation imaging assay. (**A**) Schematic illustration of luciferase complementation imaging assay upon transient expression of test genes in *N. benthamiana* leaves. Constructs of *MIF* and all three *MDL* genes were N-terminally fused to N- and C-terminal segments of firefly luciferase, respectively. The resulting plasmids were transferred into *A. tumefaciens* strain GV3101 (pmP90RK) for subsequent transformation of the plasmids into plant cells. For co-infiltration, equal volumes of each *A. tumefaciens* transformant culture were mixed and infiltrated with a syringe lacking a cannula from the abaxial side into fully expanded leaves of four- to six-week-old *N. benthamiana* plants. Imaging was done after three days of incubation following spraying the leaves with the corresponding substrate D-luciferin. (**B**) Immunoblot analysis of transient expression of luciferase complementation imaging fusion proteins in *N. benthamiana* leaves. Protein extracts of *Agrobacterium*-infiltrated *N. benthamiana* leaves were separated by SDS-PAGE and blotted onto a nitrocellulose membrane. The blot was probed with an anti-luciferase primary antibody and a secondary antibody coupled to horseradish peroxidase. Chemiluminescence detection of antigen-antibody complexes was performed with SuperSignal™ West Femto Western substrate. As a loading control, nitrocellulose membranes were stained in Ponceau S solution. Expected molecular masses are ∼57 kDa for the nLUC-MIF/MDL fusion proteins and ∼29 kDa for the cLUC-MIF/MDL fusion proteins.

**Supplemental Figure 7.**
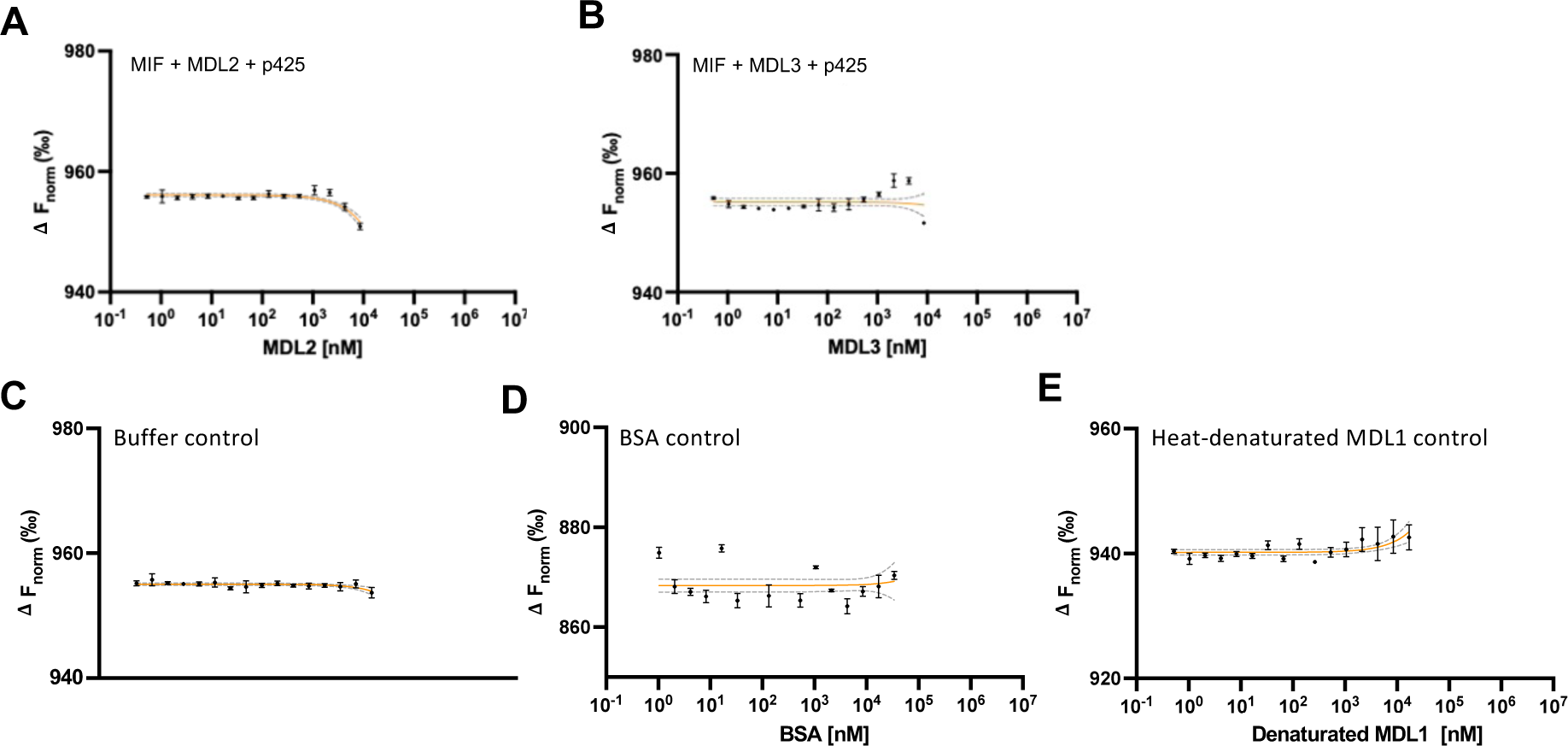
Effect of p425 with MIF-MDL2 and MIF-MDL3 and control experiments for microscale thermophoresis (MST). RED-NHS-MIF was tested for interaction in MST with different controls. Settings and buffer conditions were the same as for the MIF-MDL experiments (20 mM sodium phosphate buffer, pH 7.2, containing 0.2% Tween-20). Values shown represent means ± SD as obtained from at least three independent experiments. Data analysis and KD-fitting was performed using NanoTemper MOcontrol software, visualization was done *via* non-linear fitting using Graphpad Prism. MIF and p425 at different concentrations of (**A**) MDL2 and (B) MDL3, Controls with (**C**) Buffer control, (**D**) bovine serum albumin (BSA) as an unrelated control protein and (**E**) heat-denatured MDL1 as a negative control protein.

**Supplemental Figure 8.**
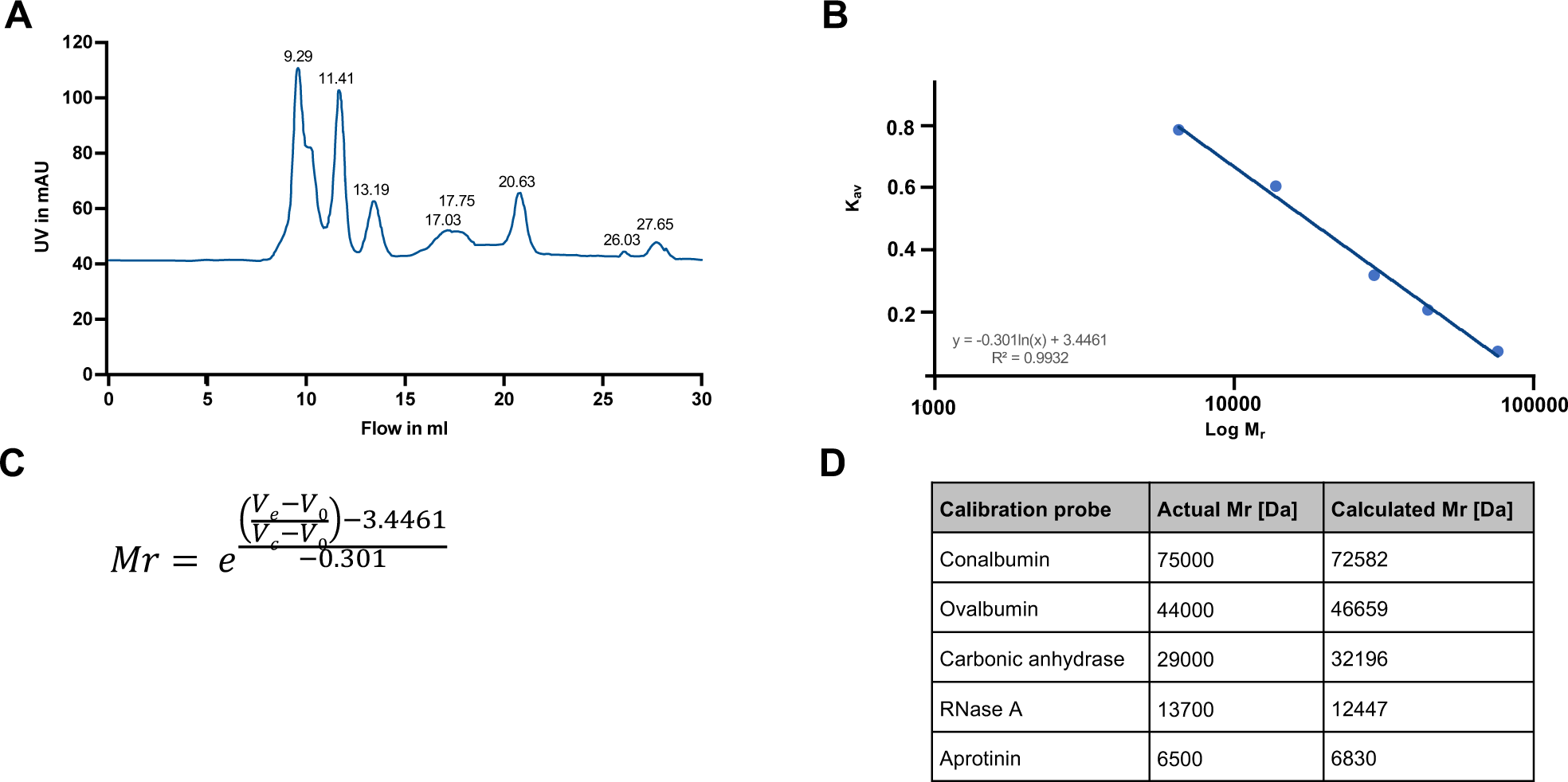
Establishing a standard curve and standard equation for the Superdex 75 10/300 SEC column. The GE Healthcare Gel Filtration Calibration Kit was used to establish a standard curve and equation for the following conditions: 20 mM sodium phosphate buffer including 20 mM sodium chloride, pH 7.2, flow rate 0.5 mL/min. (**A**) Standard proteins with known molecular masses were prepared, mixed according to manufacturer’s instructions and run over the column under the aforementioned conditions. The chromatogram shows the elution profile of the standard proteins with their corresponding elution volumes. (**B**) Standard curve generated from the known molecular mass and the observed elution volume for each of the test proteins. Notice the logarithmic x-axis. (**C**) Standard equation to calculate the molecular mass (Mr) of a protein according to its elution volume (Ve). V0 = column volume, e = Euler’s number. (**D**) Comparison of the known molecular masses of test proteins to their calculated mass based on their elution volumes (Ve) and the standard equation shown in **C.**

## Notes

### Competing Interest Statement

The authors have declared no competing interest.

